# Phenotypic and Functional Alterations in Peripheral Blood Mononuclear Cell-Derived Microglia in a Primate Model of Chronic Alcohol Consumption

**DOI:** 10.1101/2025.02.05.636708

**Authors:** Hami Hemati, Madison B. Blanton, Heather E. True, Jude Koura, Rupak Khadka, Kathleen A. Grant, Ilhem Messaoudi

## Abstract

Alcohol-induced dysregulation of microglial activity is associated with neuroinflammation, cognitive decline, heightened risk for neurodegenerative diseases, alcohol dependence, and escalation of alcohol drinking. Given the challenge of longitudinally sampling primary microglia, we optimized an *in vitro* method to differentiate peripheral blood mononuclear cells (PBMC) from non-human primates (NHP) into microglia-like cells (induced-microglia; iMGL). The iMGLs displayed transcriptional profiles distinct from those of monocyte progenitors and closely resembling those of primary microglia. Notably, morphological features showed that differentiated iMGLs derived from NHPs with chronic alcohol consumption (CAC) possessed a more mature-like microglial morphology. Additionally, dysregulation in key inflammatory and regulatory markers alongside increased baseline phagocytic activity was observed in CAC-derived IMGLs in the resting state. Phenotypic and functional assessments following LPS stimulation indicated the presence of an immune-tolerant phenotype and enrichment of a CD86^+^ hyper-inflammatory subpopulation in iMGLs derived from ethanol-exposed animals. Collectively, these findings demonstrate that *in vitro* differentiation of PBMC offers a minimally invasive approach to studying the impact of CAC on microglial function revealing that CAC reshapes both functional and transcriptional profiles of microglia.

## INTRODUCTION

In 2020, among all the people with Substance Use Disorder, 71% struggled with alcohol use^1^. Additionally, 10.5% of people ages 12 and older (29.5 million people) suffered from Alcohol Use Disorder (AUD) in 2022, according to the National Survey on Drug Use and Health in the United States^2^. AUD is a disorder mediated by psychosocial, biochemical, and genetic factors and correlates with heightened hospitalization rates^3^, intensive care unit admissions^4^, and increased incidence of multiple diseases^5^, including neurological diseases^6–10^. Data from several studies indicate that both acute and chronic alcohol consumption (CAC) dysregulate the central nervous system (CNS)-resident cells^11^, leading to behavioral changes, including escalated ethanol use, compulsive seeking, dependence^12–14^, and the risk of post-abstinence drinking^11^. These impacts are likely regulated through changes in the activity of transcription factors and alterations in epigenetic regulation, alternative splicing, protein translation, post-translational modifications, and changes in neuromodulators and neural circuits^15^ that together disrupt brain homeostasis by affecting various brain-resident cell types, including microglia^16^.

Microglia- the brain’s resident immune cells- are fetal yolk sac-derived macrophages that regulate brain development and tissue homeostasis. They maintain neuronal networks by regulating neuronal repair, survival and activity, synapse pruning, and myelination^17–19^. Therefore, disturbances in microglial function might lead to altered cognitive processes^20^, as well as learning and memory deficits^21^. Alcohol consumption-induced dysregulation of microglial responses may enhance alcohol craving and increase the risk of post-abstinence drinking^22^ and alcohol dependence^23, 24^. Studies in rodent models of alcohol consumption have shown that depletion of microglia or blockade of microglia activation decreased alcohol self-administration, reduced withdrawal-induced anxiety and post-abstinence drinking, and inhibited proinflammatory gene expression after acute withdrawal from binge alcohol^24–26^.

However, these findings are mostly derived from rodent models of CAC that do not always closely recapitulate human alcohol drinking and its consequences. Since obtaining human microglia for comprehensive experiments is unfeasible, alternative methods to study the interplay between AUD and microglia dysfunction have been developed. These include using immortalized cell lines^27^, embryonic stem cells (ESC)- and induced pluripotent stem cells (iPSCs)-derived microglia in a 2D or 3D organoid culture^28^ or a co-culture system with neurons^29, 30^. However, these models have several limitations^31, 32^. For instance, pluripotent or endothelial stem cells are developmentally far from microglia. In addition, some of the commercially available approaches are lengthy, taking 30 to 70 days^33^.

To address these limitations, we optimized a method to differentiate PBMC into microglia (induced microglia: iMGLs) based on a previously reported method^34–36^. It was shown that iMGLs are a functionally valid model for studying microglia characteristics^37, 38^. This approach allowed us to derive microglia from PBMC collected from a rhesus macaque model of voluntary ethanol self-administration in the absence and after 12 months of voluntary ethanol self-administration. This method offers a robust platform for evaluating biological outcomes in an outbred species while overcoming common confounders of clinical studies such as variation in diet, housing, polysubstance use, and infections. Our results show profound changes in morphological profiles and functional characteristics, including cytokine secretion and phagocytosis, as well as transcriptome profiles at resting and following stimulation with LPS.

## MATERIALS AND METHODS

### Animal study and PBMC and brain samples

We used cryopreserved PBMC obtained from rhesus macaques that engaged in voluntary ethanol consumption as described in^39–41^. Briefly, the animals were introduced to a 4% w/v ethanol solution during a 90-day induction period, followed by the option to drink a 4% ethanol w/v solution or water for 22 hours/day for one year^42, 43^. This setup yields stable drinking patterns, with approximately 40% of these animals developing into heavy or very heavy drinkers. The PBMC samples used in this study were obtained from either controls or macaques after 12 months of heavy ethanol consumption (CAC: chronic alcohol consumption); PBMC control group: n=11 (7 Female, 4 Male), PBMC CAC group: n=11 (6 female, 5 male). The brain samples were obtained from animals after 6 months of CAC: n=8 (4 female, 4 male). The samples were acquired through the Monkey Alcohol Tissue Research Resource (www.matrr.com). This study was approved by the Oregon National Primate Research Center (ONPRC) Institutional Animal Care and Use Committee.

### Primary microglia isolation

Primary microglia were isolated from biopsies (0.23±0.08 gram) obtained from three brain regions: SFO (subfornical organ; a region in the circumventricular organ (CVO)), area 46 (a subdivision of the prefrontal cortex), and insula (a region of the cerebral cortex) from animals after 6 months of CAC: n=8 (4 female, 4 male). The samples were shipped in Hibernate-A medium (Gibco) and processed immediately upon arrival within 24 hours, as previously described^44, 45^. Briefly, the tissues were minced with a scalpel blade and then incubated in digestion media (HBSS without Calcium (Ca^2+^) and magnesium (Mg^2+^), 5% phosphate-buffered saline (PBS), 10µM HEPES, 2mg/mL Collagenase A, 28U/mL DNAse 1, 1% Penicillin-Streptomycin) at a ratio of 5 ml digestion buffer for 0.5-1g/tissue in a 37°C water-bath for 35 minutes. Every 5 minutes, the sample was removed from the water bath and mixed ∼20 times by progressively reducing the aperture of the pipette tip (transfer pipette, 1000uL pipette tip, 200uL pipette tip 1X, 10uL pipette tip 2X). The sample was then filtered through a 70 mm nylon filter, washed with HBSS containing Ca^2+^/ Mg^2+^, and centrifuged at 300 g for 10 minutes at 4°C, with maximum acceleration and break set to half speed. The pellet was then resuspended in 20 mL of DMEM (Dulbecco’s Modified Eagle Medium) without phenol red, supplemented with 10% FBS and 1% Penicillin-Streptomycin. Using a 10 mL serological pipette set on slow release, 10 mL of Percoll was directly added to the sample dropwise and spun for 30 minutes at 4000 rpm with maximum acceleration and break at half speed at 4°C. After centrifugation, the sample was separated into three layers: myelin on top, glial cells, erythrocytes towards the bottom, and Percoll on the bottom. The glial cells were collected using a transfer pipette and washed twice using 25mL of DMEM supplemented with 10% FBS, 1% Penicillin-Streptomycin, and 25 mM HEPES Buffer, and centrifuged for 10 minutes at 1500 rpm at 4°C for the first wash and then centrifuged at 10 minutes at 1200 rpm at 4°C for the second wash. Cells were then counted, frozen in a 90-10% FBS/Dimethylsulfoxide (DMSO) solution, and stored until samples could be processed as a batch.

### PBMC-derived microglia culture/stimulation

We modified a previously reported method^34–36^ for generating microglia from PBMCs. Briefly, 24-well plates were coated with Poly D-lysine at a final concentration of 50 µg/ml overnight at 37°C in a CO2 incubator, shielded from light. The following day, the Poly D-lysine was removed, and the wells were washed three times with cell culture-grade water (Gibco). The plates were then dried. PBMCs were thawed in RPMI-1640 (Roswell Park Memorial Institute Medium) supplemented with 10% FBS, 1% Penicillin-Streptomycin, and 1% DNase. The cells were counted and mixed with the seeding media (RPMI 1640 containing 10% FBS and 1% Penicillin-Streptomycin). 1X10^6^ cells were seeded in each well with 500 µl of the media. The plates were then incubated for 16 hours at 37°C in the incubator. Subsequently, the media was replaced with induction media, which consisted of RPMI 1640 supplemented with 1% GlutMAX, human recombinant IL-34 (10 µg/ml), human recombinant GM-CSF (5 µg/ml), and 1% Penicillin-Streptomycin. The plates were then incubated for an additional 13 days. The media was then replaced with induction media. After 24 hours, the cells were collected for RNA extraction or stimulated with Lipopolysaccharides (LPS) at a final concentration of 1 µg/ml for 4 hours.

### RNA isolation and bulk RNA-seq library preparation

Differentiated cells were lysed using QIAzol. Total RNA was then isolated using Qiagen’s mRNeasy kit following the manufacturer’s instructions, and RNA integrity and concentration were determined using Agilent’s 2100 Bioanalyzer. Libraries were prepared with New England Biolabs NEBnext Ultra II Directional RNA Library Prep Kit. Briefly, NEBNext® rRNA Depletion Kit v2 (Human, Mouse, Rat) was used for rRNA depletion. The remaining mRNA was then fragmented, converted to cDNA, and ligated to adaptors. ∼300bp segments were selected using NEBNext Sample Purification Beads, amplified via PCR, and multiplexed. Library quality was assessed for concentration, size, and quality on Agilent’s 2100 Bioanalyzer. Samples were sequenced on the Novaseq X to an average depth of 20 million 100bp reads/sample.

### RNA-Seq data analysis

Bulk RNA-seq data of monocytes obtained from macaques after 12 months of CAC (3 female) were retrieved from our previous publication^46^. Microglia libraries and monocytes sequences were quality assessed using Fast QC (https://www.bioinformatics.babraham.ac.uk/projects/fastqc/), and TrimGalore was utilized to remove ligated adaptors and remove low-quality base calls, ensuring a final PHRED score of 30 and a minimum length of 70bp. HITSAT^47^ was utilized to align reads to the *Macaca mulatta* (Mmul_10) reference genome based upon gene annotations available on ENSEMBL Biomart^48^, and the GenomicFeatures package in R was utilized to determine alignment counts. Aligned reads were normalized by transcripts per million (TPM). Differentially expressed genes (DEGs) were defined as FDR ≤0.05, and a log fold change of ±1 using edgeR^49, 50^. Functional enrichment of DEGs was completed using Metascape^51^ and Gene Set Enrichment Analysis (GSEA)^52^.

### Immunofluorescence staining

Cells were seeded as described above in 24-well plates on coverslips. On day 15, coverslips were carefully transferred to a new plate and fixed with 4% paraformaldehyde (PFA) for 15 minutes at room temperature. Following fixation, cells were washed three times with phosphate-buffered saline (PBS) for five minutes each. Cells were then permeabilized by incubating coverslips in PBS with 0.2% Triton X-100 for 20 minutes on a rocker. After two additional PBS washes, blocking was performed in 5% bovine serum albumin (BSA) in PBS containing 0.1% Triton X-100 for 2 hours at room temperature. Cells were then incubated with primary antibodies diluted in 1% BSA-PBS containing 0.1% Triton X-100. Primary antibodies IBA1 (1;1000, FUJIFILM Wako Pure Chemical, 019-19741) or TMEM119 (1:1000) incubation was conducted overnight at 4°C in a humidified chamber. After three washes with blocking buffer (1% BSA-PBS) for 10 minutes each, secondary antibody Goat anti-Rabbit IgG-Alexa Fluor™ Plus 594-Red (Invitrogen) was applied at appropriate dilutions (0.5 µg/ml) in 1% BSA-PBS and incubated for 1 hour at room temperature in the dark. Coverslips were then washed three times with PBS to remove unbound antibodies. DAPI (1 μg/mL) was applied for 15 minutes to stain nuclei, followed by two PBS washes. Coverslips were then mounted onto glass slides using mounting media (Prolong Gold) and sealed with clear nail polish. Slides were allowed to dry overnight and stored at 4°C until imaging.

### Morphology analysis

Images were captured at 40X magnification with a 1 µm step size using z-stack acquisition. ImageJ software was used to identify pixels of maximum intensity within each z-stack. The 8-bit images were further processed with the ImageJ MicrogliaMorphology macro and its corresponding R package^53^. Briefly, an automatic local or global threshold was applied to each image, followed by single-cell microglia segmentation using optimal threshold settings. Skeletonization of single cells was performed, and fractal analysis was conducted with the FracLac plugin. Images were color-coded using the ColorByCluster feature. The 27 morphological features were extracted and plotted to reveal group-specific morphological differences. These features were further analyzed using the MicrogliaMorphologyR package, employing Principal Component Analysis (PCA) and clustering analysis to examine cell heterogeneity and identify morphologies unique to each group.

### Spectral flowcytometry and dimensionality reduction analysis

Microglia phenotype was established using the following markers; CD14, CD68, HLA-DR, CD11b, IBA1, CX3CR1, PU.1, P2RY12, TMEM119, CD16, CD40, CD86, CD163, CD206, TREM2, IRF8, and CD115 (GSF1R)^54, 55^ using the spectral analyzer CyTek Arora. Briefly, iMGLs were collected, washed, and incubated with a cocktail including cell surface antibodies in presence of Brilliant Stain buffer (50µl/sample, BD Biosciences), TruStain FcX Fc Receptor Blocking Solution (5µl/sample, Biolegend), and True-Stain Monocyte Blocker (5µl/sample, Biolegend) in 5 ml Tubes for 30 minutes at 4°C in the dark. The cells were then washed with PBS and fixed with Tonbo permeabilization solution for 2 hours at 4°C in the dark. Cells were then washed with Tonbo Perm buffer and stained with intracellular antibodies overnight at 4°C in the dark. Subsequently, the cells were washed and resuspended in stain buffer (PBS+ 2% FBS+ 1 mM Ethylenediaminetetraacetic acid (EDTA)). Samples were acquired on CyTek Aurora flow cytometer (5-laser; 355 nm, 405 nm, 488 nm, 561 nm, and 640 nm) using SpectroFlo Software v2.2.0.2. The cells were unmixed using stained beads with the autofluorescence extraction option enabled. The unmixed FCS files were used for further data analysis on Flowjo 10.10.0. Briefly, undesired events were removed by FlowAI 2.3.2^56^, and batches were corrected using cyCombine 1.0.2^57^. For supervised gating, the CD45^+^CD11b^+^ population was identified as the iMGLs parent population. Cells expressing P2RY12^+^TMEM119^+^ were designated as iMGLs. The expression of surface and intracellular markers on iMGLs was then assessed. The CD45^low-to-high^CD11b^low-to-high^ population was selected for unsupervised analysis to capture all intermediate populations. This population was exported for downstream dimensionality reduction using UMAP 4.1.1 (Uniform Manifold Approximation and Projection for Dimension Reduction)^58^ and clustering by FlowSOM 4.1.0^59^. PHATE (Potential of Heat-diffusion for Affinity-based Trajectory Embedding)^60^ was used to visualize cluster relations.

### Phagocytosis

pHrodo Red *E. Coli* BioParticles (Invitrogen) was utilized to survey the phagocytic capacity of microglia. On day 15, the cells were incubated with pHrodo Red *E. Coli* BioParticles for 4 hours. Cells were then stained for CD45, CD11b, TMEM119, and P2RY12 and acquired on the Attune NxT Cytometer (ThermoFisher Scientific). The data were analyzed using FlowJo 10.10.0.

### Luminex

The supernatants were collected from differentiated cells before or after LPS stimulation on day 15, centrifuged for 10 min at 10000 rpm to remove cell debris, and immediately stored at -80. Immune mediator production was determined using a Millipore NHP-validated 10 plex kit (MCP1, MIP-1β, IL-6, IL-8, IFN-γ, IL-10, IL-1β, IL4, TNF-α, MIP-1α). Data was acquired using a MAGPIX instrument and analyzed using a 5-parameter logistic regression with the xPONENT™ software.

### Statistical analyses

We conducted a normality assessment using the Shapiro-Wilk test (alpha=0.05) and identified outliers through ROUT analysis (Q=0.1%). If the dataset showed adherence to a normal distribution, group differences were assessed using either paired or unpaired t-tests with Welch’s correction. Multiple comparisons were addressed using the Holm-Sidak test, which adjusted the family-wise significance and confidence level to 0.05. In cases where the data did not meet Gaussian assumptions, group comparisons were conducted using the Mann-Whitney test.

## RESULTS

### Peripheral monocyte-derived microglia serve as a model for studying microglia in rhesus macaques

The iMGLs were analyzed for their phenotypic, morphological, and functional characteristics **(Fig. 1A)**. We first undertook a series of assays to confirm that the differentiated microglia exhibit phenotypic and functional characteristics that closely resemble those of primary microglia. The differentiation efficiency was assessed by measuring the expression of microglia canonical markers throughout the differentiation process. To identify microglia progenitors, we used CD14 as a common marker of monocytes and microglia and CX3CR1, which is expected to increase in expression throughout the maturation of microglia^61^. While less than 5% of CD14^+^CX3CR1^+^ progenitor cells were P2RY12^+^ TMEM119^+^ (iMGL) on day 1, this percentage reached 100% on day 15 **(Fig. 1B)**. This was accompanied by the gradual increase in the expression of the microglia marker TMEM119 **(Supp Fig. 1A)**. Additionally, the percentage of CD45^+^CD11b^+^ cells exceeded 50% on day 15, with over 92% of these cells being P2RY12^+^TMEM119^+^ (**Supp. Fig. 1B**). The bright-field and confocal imaging showed enhanced expression of microglia canonical markers IBA1 (**Fig. 1C**) and TMEM119 in differentiated microglia (**Supp. Fig. 1C**). Finally, the functionality of the iMGLs was validated by stimulating them with LPS, which resulted in TNF⍺ production **(Supp. Fig. 1D)**.

**Figure 1.**
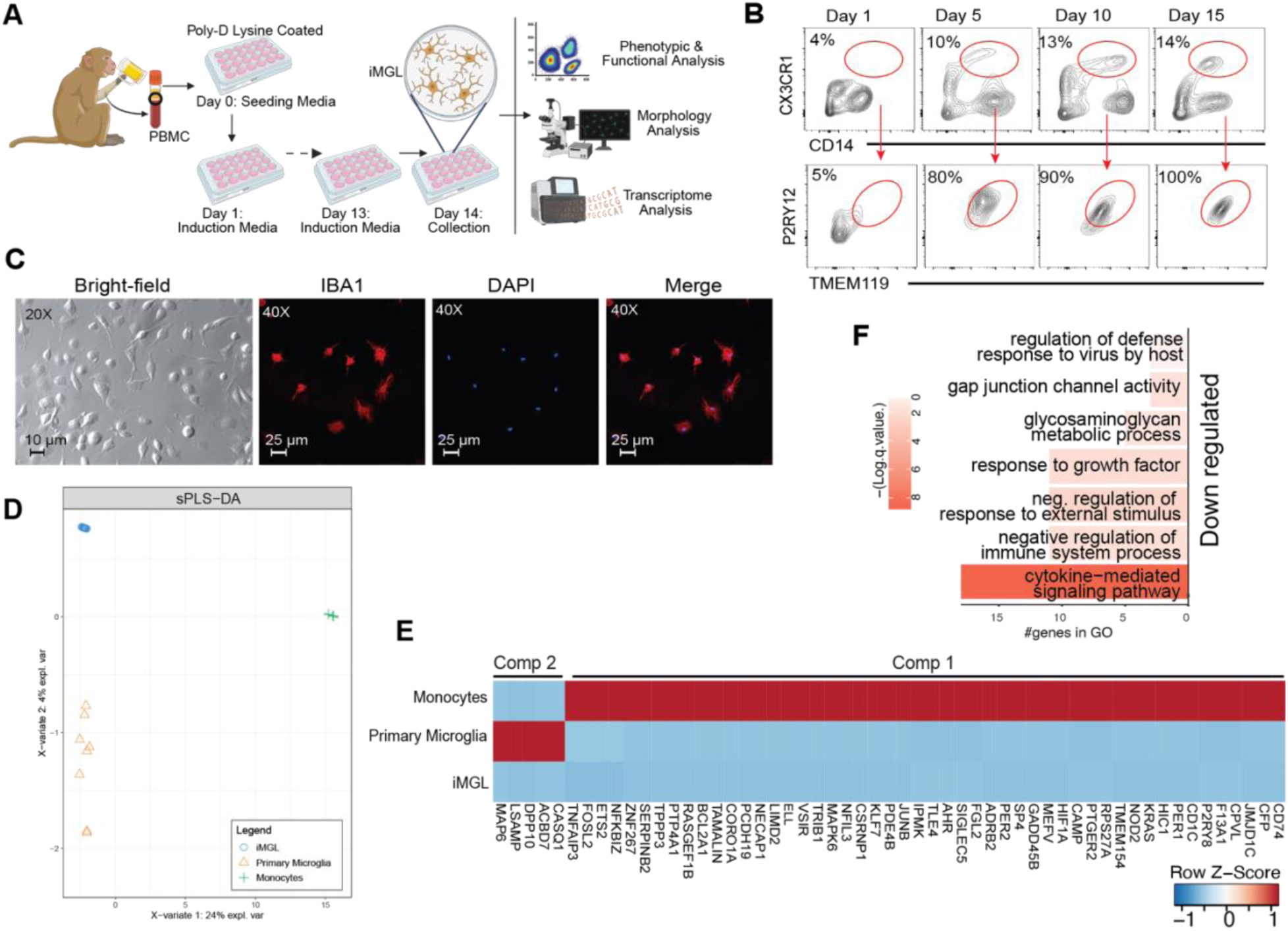
Rhesus macaques PBMC-derived microglia exhibited transcriptional similarities to primary microglia. **A)** Experimental design. **B)** Frequency of CD14^+^CX3CR1^+^ cells and P2RY12^+^TMEM119^+^ as determined by flow cytometry. **C)** Bright-field imaging (20X) and confocal imaging (40X) of IBA1 staining. **D)** (s)PLS-DA analysis of the transcriptome profile of primary microglia, iMGLs, and monocytes. **E)** Heatmap depicts the expression of genes driving the differences observed along Components 1 and 2. **F)** Downregulated genes in iMGLs compared to primary microglia were enriched for immune signaling.

Next, we assessed the extent to which the transcriptome of iMGLs diverged from that of their monocyte progenitors and approached that of primary microglia isolated from rhesus macaque through RNA sequencing. The primary microglia were isolated from the circumventricular organ (CVO), the prefrontal cortex, and the insula, as these areas play a critical role in behavioral regulation^62^ and decision-making and have been implicated in the control of alcohol consumption^63, 64^. (s)PLS-DA analysis showed that the transcriptional profile of iMGLs is distinct from that of monocytes and closely related to that of primary microglia **(Fig. 1D)**. Enrichment of genes in components 1, which delineated monocytes from both iMGLs and primary microglia were involved in inflammatory processes (ex. *TNFAIP3, NFKBIZ, NOD2. HIF1A, JUNB*). On the other hand, genes in component 2 played a role in neuronal health and signaling (e.g., *LSAMP*, *MAP6*, and *DPP10*) **(Fig. 1E & Supp. Fig.2 A)**.

We then directly compared gene expression between primary microglia and iMGLs. The few differentially expressed genes (DEGs) upregulated in iMGLs relative to primary microglia did not significantly enrich to any GO terms. However, among them, we observed genes involved in lipid metabolism (*AQP10*, *CIDEC*, *FGF21*), neurological signaling (*RLN3*, *KCNK17*), a component of the mitochondrial electron transport chain (*COX6B2*). On the other hand, DEGs downregulated in iMGLs relative to primary microglia enriched to terms associated with regulatory (ex. *GJA, SRC*) and inflammatory pathways (ex. *IL1B, IL12, TNF*), indicating a potentially less mature state of the iMGLs **(Fig. 1F & Supp. Fig.2B-C)**. Overall, this observation supports that iMGLs are transcriptionally close to primary microglia and, therefore, are an appropriate *in vitro* model to study primary microglia.

### Chronic alcohol consumption alters the morphology of iMGLs

Chronic alcohol consumption has the potential to disrupt the brain microenvironment through sustained activation of microglia^65^. Activated primary microglia transform from a highly ramified shape to a hypertrophic and amoeboid-like shape with increased phagocytic and proliferation capacities^66–69^. To evaluate the impact of CAC on the morphology of the iMGLs, confocal images were processed using the ImageJ macro MicrogliaMorphology and its R package^53^, with slight modifications to adapt it to *in vitro* microglia culture. The extracted 27 morphology features were plotted to identify the overall differences between the groups and further evaluated using dimensionality reduction and clustering analysis to capture the heterogeneity among the cells to define morphologies specific to each group (**Fig. 2A**). This analysis showed that CAC-derived iMGLs possessed an overall lower “circularity” with the enhanced “perimeter” that indicates an overall decrease in the presence of ameboid morphology among CAC-derived iMGLs (**Fig. 2B**).

**Figure 2.**
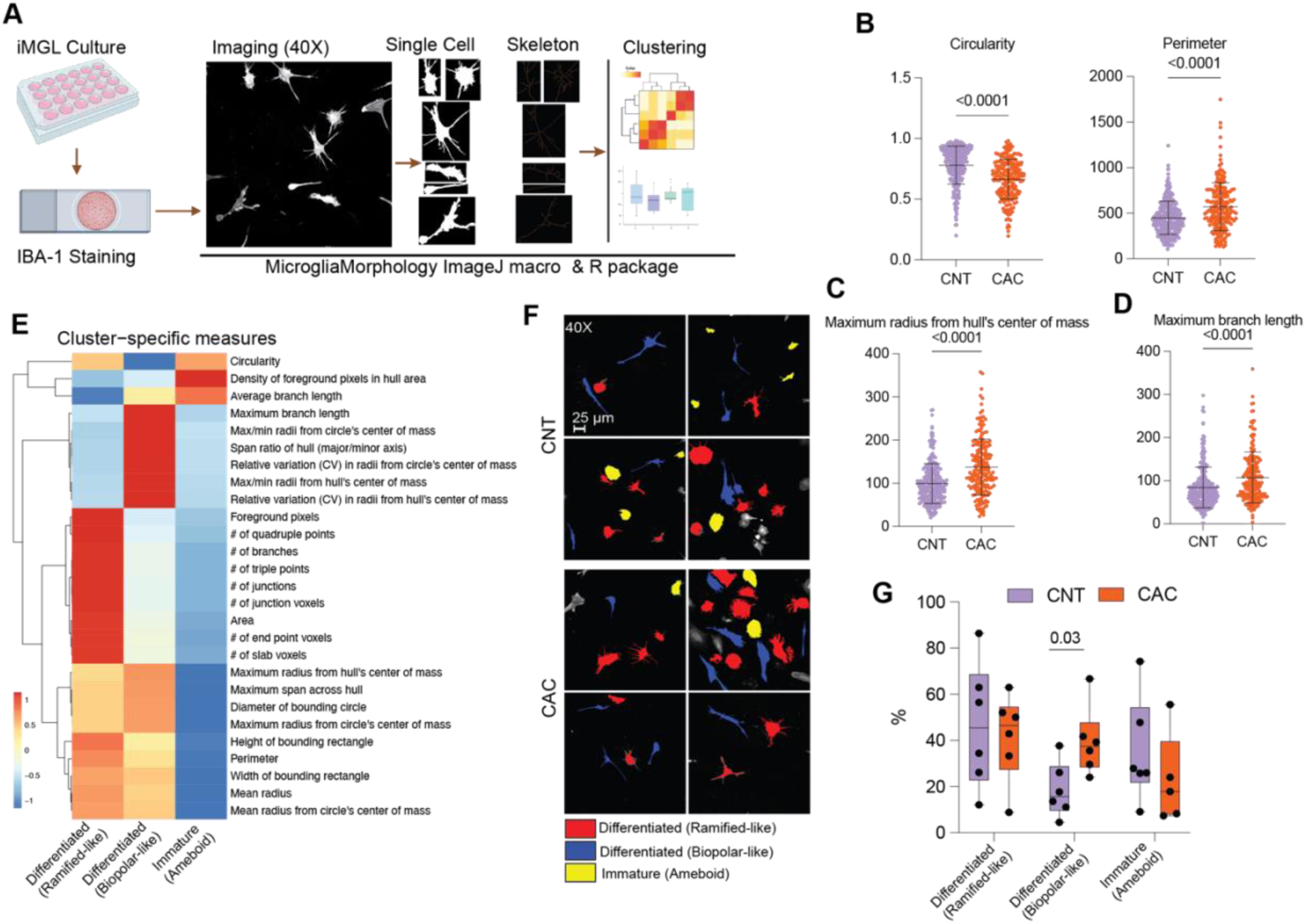
CAC-derived PBMCs differentiated into iMGLs with a more mature-like morphology. **A**) Experimental design for morphology analysis. Confocal images were used to determine the morphological features; **B**) “circularity” and “perimeter”, **C**) “Maximum radius from hull’s center of mass”, and **D**) “Maximum branch length”. **E**) Heatmap of cluster-specific measures on average; the measures were scaled across clusters. **F**) The individual cells were spatially registered back to the original images and visually annotated based on morphological classes using the ColorByCluster feature. **G**) Frequency of three identified morphological clusters. For B-D, an unpaired Mann-Whitney t-test was used to compare the features (p≤0.05). E and F were generated using the default setting of the MicrogliaMorphologyR package unless otherwise specified. See Supp. Fig.3 for additional details.

We also detected a significant increase in features describing area and territory span, including enhanced “Maximum radius from hull’s center of mass” **(Fig. 2C**). In addition, significant increased “Maximum span across hull”, “Max/min radii from hull’s center of mass”, “Width of bounding rectangle”, “Height of bounding rectangle”, Maximum radius from circle’s center of mass, “Diameter of bounding circle”, and “Mean radius” in CAC-derived iMGLs indicated that their overall size was larger than that of controls (p≤0.05) (**Supp. Fig. 3A**), suggestive of enhanced differentiation with CAC.

In addition, features describing cell shape including “Span ratio of hull (major/minor axis)”, “Relative variation (CV) in radii from hull’s center of mass”, and “Relative variation (CV) in radii from circle’s center of mass” (**Supp. Fig. 3B**), and the branch feature “Maximum branch length” **(Fig. 2D**) were significantly enhanced in CAC-derived iMGLs (p≤0.05). In contrast, the “Density of foreground pixels in the hull area” decreased in CAC-derived iMGLs (**Supp. Fig. 3C).** This feature describes the occupancy of cells within their territory that represent soma and/or branch thickness^53^, which is correlated with other observations showing the presence of branches in the hull area and fewer ameboid-like iMGLs in the CAC-group.

While these comparisons offer valuable data regarding the general distinctions among these groups, they do not fully encapsulate morphological heterogeneity. Therefore, we conducted a more comprehensive evaluation by employing dimensionality reduction and clustering analysis using MicrogliaMorphologyR 1.1^53^. The heatmap of correlations between principal components (PCs) and features revealed that PC1 is negatively associated with features reflecting branching complexity and territory span. PC2 captured variability driven by cell shape, displaying a negative correlation with “circularity” and a positive correlation with “maximum/minimum radii from the center”, the “span ratio of the hull”, and branching homogeneity (“relative variation (CV) from the center of mass”). Furthermore, PC3 showed a positive correlation with “average branch length”, and PC4 displayed a negative correlation with the “density of foreground pixels in the hull area” indicating cells that are less compact and more spread out or have extensions (**Supp. Fig. 3D).** Plotting PC1 and PC2 revealed notable distinctions between the control and CAC-derived iMGLs (**Supp. Fig. 3E).**

Subsequently, a *fuzzy K*-means clustering analysis was conducted to evaluate the first four principal components, which generated three clusters (**Supp. Fig. 3F).** By scaling the average values for all morphology measures across clusters and visual confirmation using the ColorByCluster feature in MicrogliaMorphology, we were able to define each cluster based on morphology measures in comparison to the other clusters **(Fig. 2E&F).** Cluster 1 had the greatest branching complexity, evidenced by a high number of “slab voxels”, “endpoint voxels”, “quadruple points”, “junctions”, “junction voxels”, “branches”, and “triple points”. Cluster 1 also had a high territory span and “circularity”. Together, these features resemble well-differentiated cells with ramified-like morphology. Cluster 2 had the lowest “circularity” and an average territory span, and the highest “maximum branch length”, “span ratio of hull (major/minor axis)”, and “Max/min radii from hull’s center of mass” resembled well-differentiated cells with bipolar-like morphology. Cluster 3 had the lowest territory span, “high circularity,” smallest “branch lengths,” lowest branching complexity, and territory span, and the highest “Density of foreground pixels in hull area,” resembling the “Immature” cells **(Fig. 2E&F).** The comparison of the frequency of each cluster revealed a significantly enhanced bipolar-like morphology in CAC-derived iMGLs (p≤0.05) **(Fig. 2G).** Our observations indicate that PBMCs from the CAC group differentiated into iMGLs, exhibiting morphological characteristics consistent with a more mature phenotype compared to the control group.

### Chronic alcohol consumption alters the phenotype of iMGLs

The phenotype of iMGLs was assessed via spectral flow cytometry based on the expression of microglia canonical, inflammatory and regulatory markers; CD14, CD68, HLA-DR, CD11b, IBA1, CX3CR1, PU.1, P2RY12, TMEM119, CD16, CD40, CD86, CD163, CD206, TREM2, IRF8, and CD115^54, 55^. The CD45^+^CD11b^+^ population was identified as the iMGL parent population by supervised analysis. Cells expressing P2RY12^+^TMEM119^+^ were designated as iMGL to assess the expression of surface and intracellular markers. The CD45^low-to-high^CD11b^low-to-high^ population was selected for unsupervised analysis to capture intermediate populations (**Fig.3A**).

**Figure 3.**
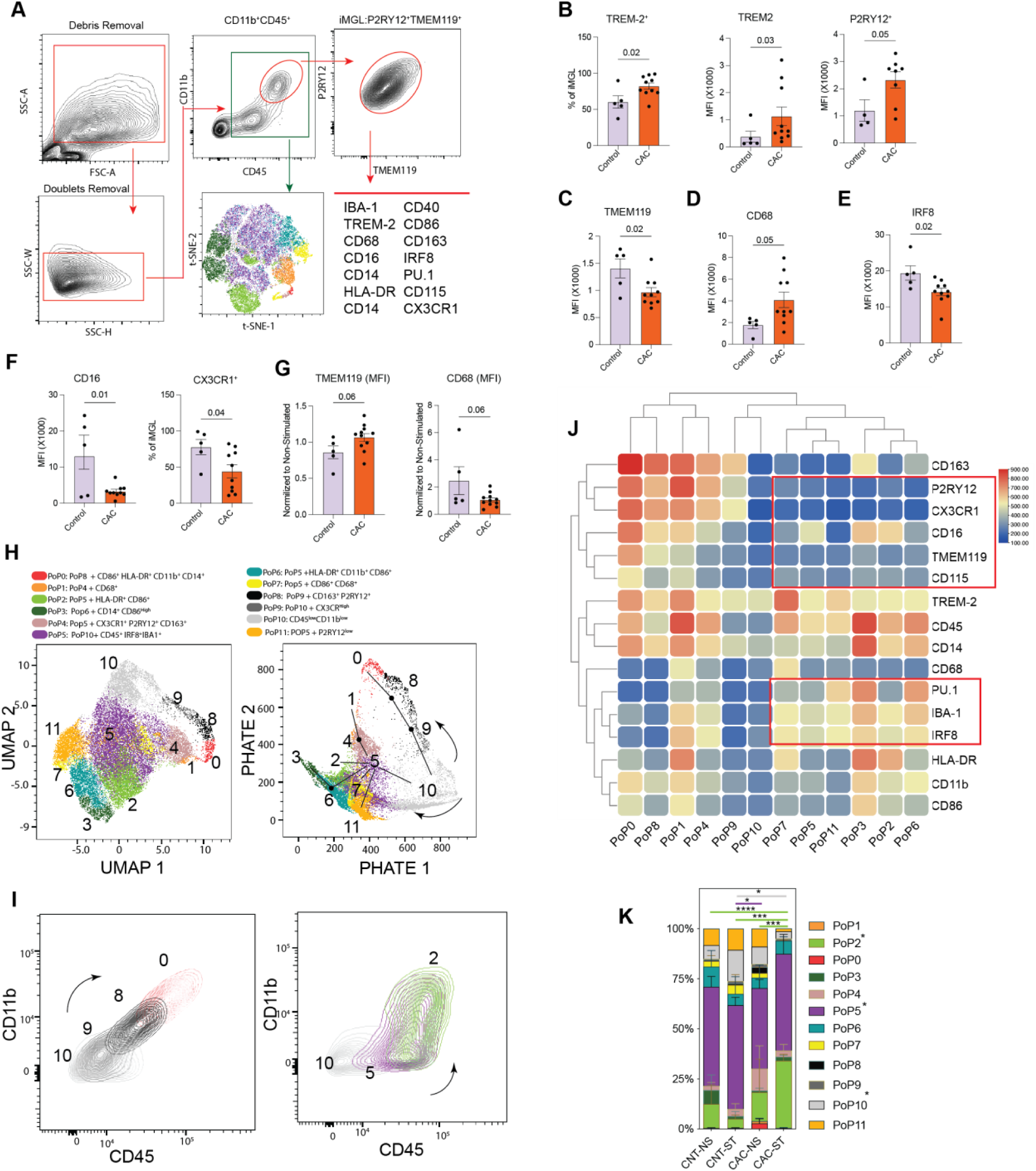
CAC skews iMGL differentiation. **A)** Full spectral flow cytometry followed by supervised and unsupervised analysis was used for phenotyping. Flow cytometry determined **B**) the frequency of TREM2+ iMGLs and expression of the level of TREM2, P2RY12, **C)** TMEM119, **D)** CD68, **E**) IRF8, and **F**) CD16 on iMGLs, as well as the frequency of CX3CR1^+^ iMGLs. **G**) Expression of TMEM119 and CD68 on iMGLs was determined following LPS stimulation. **H**) Dimension reduction was performed using UMAP, following FlowSOM to identify clusters. All the clusters were mapped on the PHATE plot. Links between clusters were drawn manually. **I**) PoPs 10, 9, 8, 0, 5, and 2 were mapped on CD11b^+^CD45^+^ population. **J**) Heatmap showing the expression level of markers across the clusters. **K**) Bar plot depicts the frequency of clusters across groups. Two group comparisons were carried out using an unpaired t-test with Welch’s correction (parametric) or Mann-Whitney (non-parametric) test. Error bars for all graphs are defined as ± standard deviation (SD).

There was no significant difference in the frequency or total number of P2RY12^+^TMEM119^+^ between the control and CAC samples (**Supp. Fig. 4 A**). Moreover, the frequency of TREM2^+^ iMGL and the MFI of TREM2 increased in the CAC-derived iMGLs (**Fig. 3B**). TREM2 plays a role in microglia proliferation through regulating metabolism^70^. This was accompanied by increased expression of cell surface markers P2RY12, which regulates microglia activation and cytokine release^71^ (**Fig. 3B**). In contrast, the expression of TMEM119 decreased with CAC (**Fig. 3C**). The precise mechanism of TMEM119 remains unclear; however, its downregulation in reactive microglia was reported^72^. Furthermore, iMGL could be categorized as CD68^high^ and CD68^low^ populations (**Supp. Fig. 4B**). Interestingly, the frequency of the CD68^high^ population and MFI of CD68 was significantly increased in the CAC-derived iMGLs (**Fig. 3D & Supp. Fig. 4B**). CD68 is a lysosomal marker that was reported to be upregulated on activated phagocytic microglia^73^. Additionally, the expression of inflammatory marker CD163 and transcription factor PU.1, which controls microglial viability and function^74^, were modestly enhanced with CAC (**Supp. Fig. 4C**). In contrast, the transcription factor IRF8 was downregulated in iMGLs derived from CAC animals (**Fig. 3E**), in line with reduced expression of CD16, CX3CR1, and CD40 (**Fig. 3F** & **Supp. Fig. 4D**). CX3CR1 expression is regulated by this transcription factor^75, 76^.

Next, we investigated iMGLs response to LPS and assessed the expression of markers in control- and CAC-derived iMGLs relative to their unstimulated counterparts. The expression of TMEM119 increased more following LPS stimulation in iMGLs derived from CAC animals compared to those derived from control iMGLs as indicated by MFI (**Fig. 3G)**. Similar observations were made for CD40 (**Supp. Fig. 4E**). In contrast, expression of CD68 was induced to a lesser extent on CAC iMGLs relative to control iMGLs following LPS stimulation (**Fig. 3G**). Importantly, the expression level of the other canonical, inflammatory, and regulatory microglia markers that are expected to be upregulated (e.g. IRF8) or downregulated (e.g. CX3CR1) were not significantly changed, following this short-term LPS stimulation.

An unsupervised analysis of the flow cytometry data using FlowSOM and Marker Enrichment Modeling (MEM) delineated 12 clusters. The Potential of Heat-diffusion for Affinity-based Trajectory Embedding (PHATE) was then used (**Fig. 3H & Supp. Fig. 4F**) to provide further insight into the developmental trajectory of the clusters, revealing two distinct lineages. PoP10, as the main iMGL progenitor cells expressing low CD45 and CD11b, gives rise to PoP5 and then PoP2, 3, 4, 6, 7, and 11 with higher IRF8, PU.1 and IBA1 expression and lower microglia markers expression. The second lineage from PoP10 to PoP0 was accompanied by enhanced expression of microglia markers P2RY12, TMEM119, CD115, and CX3CR1 and lower expression of IRF8, PU.1, and IBA1(**Fig. 3H-J & Supp. Fig. 4G**).

Comparing the frequencies of clusters among four groups-control (CNT) and CAC in non-stimulated (NS) and stimulated (ST) states-showed that PoP2 (PoP5+HLA-DR^+^CD86^+^) representing a proinflammatory subpopulation, was significantly enriched in CAC-ST samples. In contrast, the frequency of PoP5 (PoP10+CD45^+^IRF8^+^IBA1^+^) was significantly lower in CAC-NS compared to CNT-NS, and PoP10 was lower in CAC-ST compared to CNT-ST (**Fig. 3K, Supp. Fig. 4H-I**).

### Chronic alcohol consumption alters the function of iMGLs

Next, we evaluated the impact of CAC on iMGLs’ production of immune mediators in response to Toll-like receptor (TLR) ligand stimulation and their phagocytic activity. Our findings revealed a significant reduction in the production of TNF, IL-1β, IFN-γ, and MIP-1a as well as a modest decrease in levels of IL-6 and IL-8 in resting iMGLs with CAC **(Fig. 4A, Supp. Fig. 5A).** Following 4 hours LPS stimulation; we observed a significantly enhanced expression of TNF production by control iMGLs in response to LPS **Supp. Fig. 5B)**. We did not observe a significant change in the secretion of these cytokines in CAC-derived iMGLs after stimulation compared to their baseline level. Given that the baseline levels of these cytokines vary between the two groups, we evaluated the post-stimulation cytokine expression by normalizing it to their unstimulated levels, allowing us to compare responses. we observed a significant increase in the production of MIP-1b and MIP-1a by CAC-derived iMGLs compared to the control **(Fig. 4B).** However, the TNF production was not significantly different between both groups **(Supp. Fig. 5C).** Importantly, the expression of IL-1β which is downstream of LPS-TLR4 was not considerably changed **(Supp. Fig. 5D).** Additionally, the phagocytic capacity of resting iMGLs was assessed with pHrodo™ E. coli BioParticles. A modest enhancement in the uptake of *E. coli* particles was observed in iMGLs from CAC animals (**Supp. Fig. 5E & Fig. 4C**).

**Figure 4.**
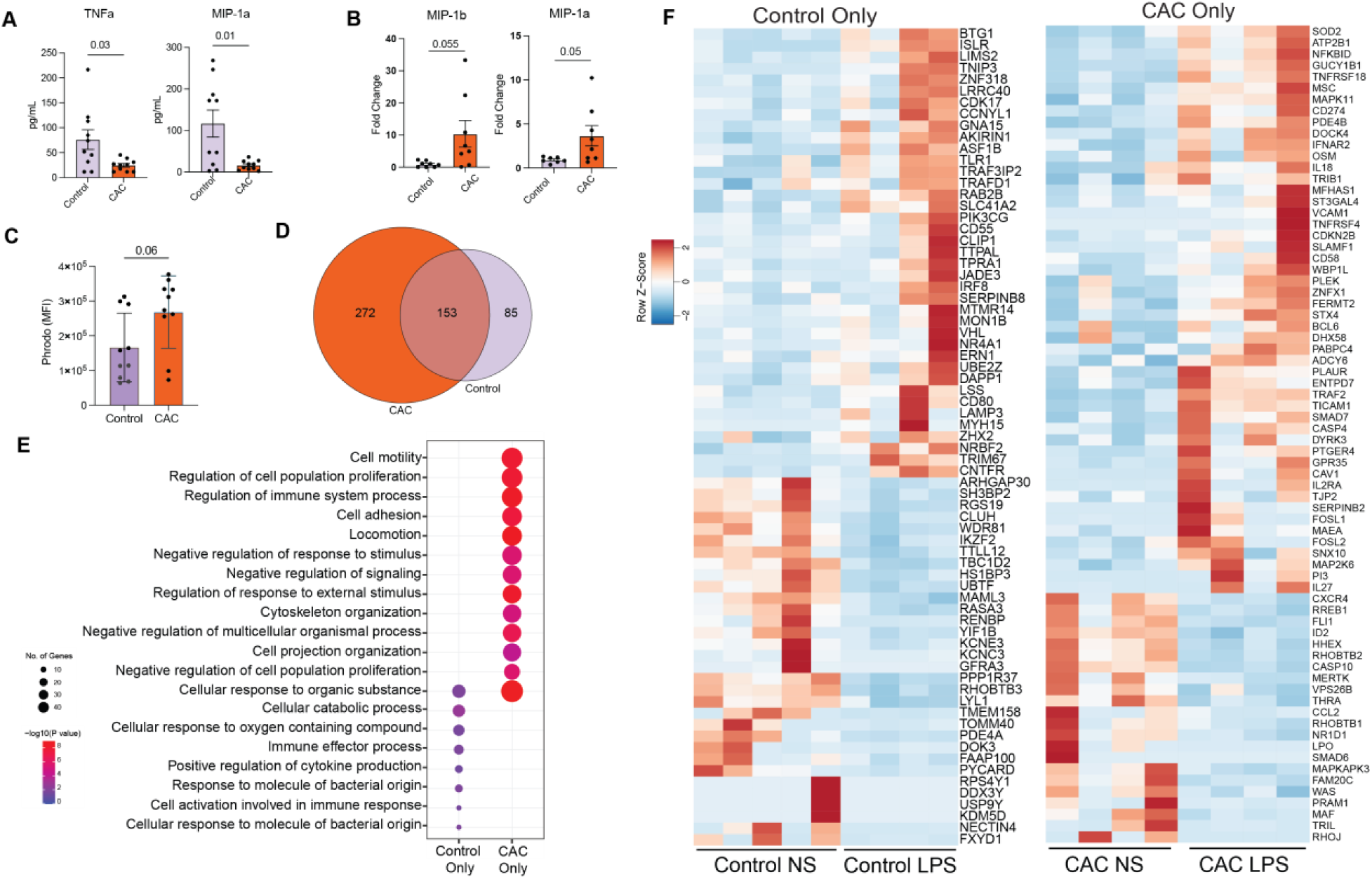
CAC-derived iMGLs exhibit dysregulated function. **A)** Production of TNF, and MIP-1α at resting and **B)** MIP-1b and MIP-1a following LPS stimulation was determined by Luminex. Expression after stimulation was normalized to unstimulated conditions. **C)** The phagocytosis ability of iMGLs at resting was assessed by pHrodo™ E. coli BioParticles. **D)** Venn diagram of DEG identified following LPS in control- and CAC-derived iMGLs **E)** Functional enrichment of DEGs unique to control or CAC groups using Metascape. Size of the bubble indicated number of DEG mapping to the GO term while color indicate FDR corrected p value. **F)** Heatmaps depicting select DEG from each group

Such variations in the secretion of immune mediators and phagocytosis activity could result from alterations in gene expression. Therefore, we analyzed the transcriptome of the resting and LPS-stimulated iMGLs using bulk RNA sequencing. While there was a large number of shared DEG in response to the LPS challenge, we also identified 272 unique genes in CAC-derived iMGLs and 85 unique DEG in controls (**Fig. 4D**). Functional enrichment analysis indicated that the DEG specific to controls enriched to processes such as “cellular catabolic process,” “positive regulation of cytokine production,” and “response to molecule of bacterial origin” (**Fig. 4E**). Notable genes that were upregulated in control-derived iMGL included *CD80*, *CD55*, *VHL*, *ERN1*, *TRIM67*, and *NR4A1* (**Fig. 4F**). On the other hand, DEGs specific to CAC-derived iMGLs enriched to processes associated with negative regulation of immune responses such as “negative regulation of cell population proliferation,” “negative regulation of response to stimulus,” and “negative regulation of signaling” which could explain the reduced response to LPS. Indeed, the expression of several negative regulators of immunological signaling, like *DUSP16*, *PIK3IP1*, *SMAD7*, *RASA2*, and *TRIB1,* was increased in iMGLs derived from the CAC group (**Fig. 4E-F**). Additional DEG unique to the CAC group mapped to GO terms associated with cell migration such as “cell motility,” “cell adhesion,” and “locomotion,” including the upregulation of genes such as *ST3GAL4*, *STX4*, and *SLAMF1* (**Fig. 4E-F**). Finally, DEGs in the CAC group also enriched to “cytoskeleton organization” and “cell projection organization” raising the possibility of morphological alterations (**Fig. 4E**).

## DISCUSSION

Alcohol Use Disorder (AUD) significantly impacts brain function, leading to cognitive deficits, such as heightened impulsivity and impaired executive decision-making^77, 78^. Among brain resident cells, microglia are crucial in governing cognitive function^20^. It was shown that the reduction in microglial activity was correlated with an 80% reduction in relapse rates, indicating that targeting microglia could mitigate cravings and support abstinence maintenance^22^. Therefore, understanding alcohol-mediated changes in microglia phenotype, morphology, and function is crucial for addressing its effects on mental health.

A significant challenge in the study of microglia is the impracticality of obtaining human primary microglial cells for research purposes. Alternative approaches include using *in vitro* cultures, such as cell lines, stem cells, or organoids, each with its limitations^31, 32^. In this study, we transdifferentiated PBMC into microglia-like cells in 2 weeks by optimizing a previously reported approach^34–36^. We found that the transcriptome profile of iMGLs is distinct from their monocyte progenitors while showing a strong similarity to primary microglia.

Microglia become activated and proliferate when triggered by brain injury, infection, or neuroinflammation. Their morphology transforms from a highly ramified shape into an amoeboid shape, facilitating migration to inflammatory sites^21^. Analysis of iMGLs morphology revealed notable enhancements in the ramifications of CAC-derived iMGLs, characterized by a reduction in circularity alongside increased size and branching complexity. Further clustering analyses indicated a significant expansion of a differentiated subpopulation exhibiting a biopolar-like morphology, indicating that CAC-derived induced iMGLs possess the capability to differentiate into more mature cell phenotypes. These morphological differences are probably associated with epigenetic differences within their progenitors^79^.

Increasing evidence calls into question the use of the oversimplified M1/M2 model when describing microglia responses to alcohol^80^. Furthermore, the traditional immunochemical markers used to identify microglial and macrophage phenotypes, such as HLA-DR and CD68, lack the specificity needed to distinguish between pro-inflammatory and anti-inflammatory states reliably^81^. Therefore, we used an extensive panel to characterize phenotypic differences between iMGLs induced by CAC. Our data revealed an elevated expression of TREM2, which plays a role in microglia proliferation and migration^70^. TREM2 was also shown to be upregulated in mice models following long-term alcohol exposure^82^. Inhibiting the TREM2 gene has been shown to improve synapse loss and the impairment of long-term potentiation (LTP). Therefore, high expression of TREM2 on iMGLs derived from the CAC group can potentially explain the deficits in synaptic plasticity and memory seen with prolonged alcohol consumption^82^.

We further observed dysregulation in various microglial regulatory markers in CAC-derived iMGLs at rest, such as elevated expression of P2RY12 and CD68, alongside reduced expression of TMEM119, IRF8, CD16, and CX3CR1 levels. P2RY12 is involved in microglial chemotaxis and sensing environmental ATP/ADP signals. It plays a crucial role in microglial responses to neuronal damage and injury^21^. CD68 is a lysosomal-associated membrane protein involved in phagocytosis and degradation of cellular debris^66^. IRF8 regulates microglial activation during inflammation^66^. CX3CR1 is essential to microglia and neuron communication^21^. The combined regulatory effect of these markers may explain the lower basal secretion of TNF and MIP-1a in CAC iMGLs compared to controls.

Microgliogenesis depends on the expression of IRF8 and PU.1, which play crucial roles in the lineage commitment and differentiation of microglia^61^, notably in developing mature CD45^+^c-kit^−^CX3CR1^+^ microglia^61^. IRF8 primarily regulates the inflammatory response of microglia, while PU.1 is essential for their survival and the expression of homeostatic genes^61, 74, 76^. We observed opposite change directions for these two transcription factors with CAC. While a modest increase in PU.1 was noted, expression of IRF8 was downregulated in line with reduced protein expression of CX3CR1, which is regulated by IRF8^75, 76^. This indicates a transition towards maintaining or restoring a homeostatic state rather than an activated or inflammatory phenotype. Therefore, while morphology analysis indicated the presence of cells with more mature-like morphology in iMGL derived from CAC, they exhibited abnormal expression of regulatory markers.

To investigate how these phenotypic changes affect function, we evaluated the impact of CAC on iMGLs cytokine expression and phagocytosis. While the control-derived iMGLs responded to LPS stimulation with enhanced TNF production, in contrast to our previous studies showing increased inflammatory response by CAC-derived circulating monocytes following LPS exposure^46, 83, 84^ and despite the significant presence of a subpopulation with enhanced CD86/HLA-DR expression and heightened expression of MIP-1α and MIP-1β following stimulation, we did not observe significant changes in other inflammatory markers or increased inflammatory cytokine production, including TNF-α and IL-1β upon stimulation in CAC-derived iMGLs.

To understand the molecular basis for the dampened response to LPS by CAC iMGLs, we examined gene expression patterns. Genes such as *NRBF2*, *PIK3CG*, *MTMR14, and IRF8,* which play critical roles in autophagy, phagocytosis, and inflammation,^85–87^ were upregulated in control-derived iMGLs in response to LPS. In contrast, the expression of several negative signaling regulators was upregulated in the CAC iMGLs. Notable examples include *DUSP16* (which negatively regulates MAP kinase)^88^, *PIK3IP1* (a negative regulator of PI3K signaling)^89^, *SMAD7* (which negatively regulates TGF-β signaling)^90^, and *RASA2* (a negative regulator of Ras signaling)^91^. Additionally, expression of genes such as *CXCR4,* which regulates microglia inflammation^92^, and CCL2, which modulates the complement system in microglia^93^, was downregulated in CAC-derived iMGLs, potentially reducing its response to short-term stimulation.

Microglia impact neuronal function and maintain neural networks through phagocytosis^94^. Alcohol consumption may impair the homeostatic function of microglia, including phagocytosis and pathogen recognition^65^. Microglial phagocytosis can be detrimental in specific circumstances, such as the excessive pruning of synapses observed in Alzheimer’s disease, while it can be beneficial in other contexts, such as the vital clearance of cellular debris following an injury. We observed enhanced phagocytosis activity in resting iMGLs with CAC, as previously reported for other induced microglia^68^. This observation aligns with the enhanced expression of CD68^73^ and TREM2^95^ as well as *MARCO*^96^ with CAC.

Although these findings underscore the utility of this approach in investigating CAC-mediated microglial impairment, it has some limitations. While we compared the iMGLs transcriptome to that of primary microglia, we have not compared their phenotype, morphology, and function. Additionally, the phagocytosis capacity and morphology need to be assessed after LPS stimulation. Our morphology analysis identified three different clusters of iMGLs; therefore, as a future direction, we will perform scRNA-seq to identify the heterogeneity among iMGLs and its correlation with their progenitor heterogeneity. Future studies should also investigate the epigenetic basis of the transcriptional and functional changes observed with CAC. Finally, a coculture system with neurons may help keep the microglia in niche-like conditions to obtain a more accurate assessment of their functions.

In summary, the results presented in this manuscript show that we successfully derived microglia-like cells from PBMC that displayed morphological and transcriptional characteristics akin to those of primary microglia. This model allowed us to explore the impact of CAC on the transcriptome, morphology, and phenotype of microglia. This model serves as a valuable tool for conducting *ex vivo* mechanistic studies to evaluate the role of microglia in substance use disorders.

## Author contributions

HH performed experiments with the help of MB and HT. RNA-seq was performed and analyzed by MB with the help of HH. HH analyzed the rest of the data and wrote the manuscript with MB and IM assistance. JK assisted in analyzing microglia morphology. KG designed and oversaw the live alcohol self-administration studies, RG helped implement the self-administration studies, oversaw the necropsy procedures, and provided the brain samples through the MATRR.com research resource. IM reviewed and edited the manuscript. All authors read the manuscript.

## Acknowledgments

We thank members of the Messaoudi laboratory for their help and feedback. We thank Dr. Delphine Malherbe for reading the manuscript and providing feedback. This work was supported by R01AA028735-04, U01AA013510-20, R24AA019431-14, P51OD011092, and F31AA031600-01A1.

## Data and code availability

Additional information and requests for resources should be directed to Dr. Ilhem Messaoudi. The datasets supporting the conclusions of this article are available on NCBI’s Sequence Read Archive (SRA# pending).

---

**Supp. Fig.1.**
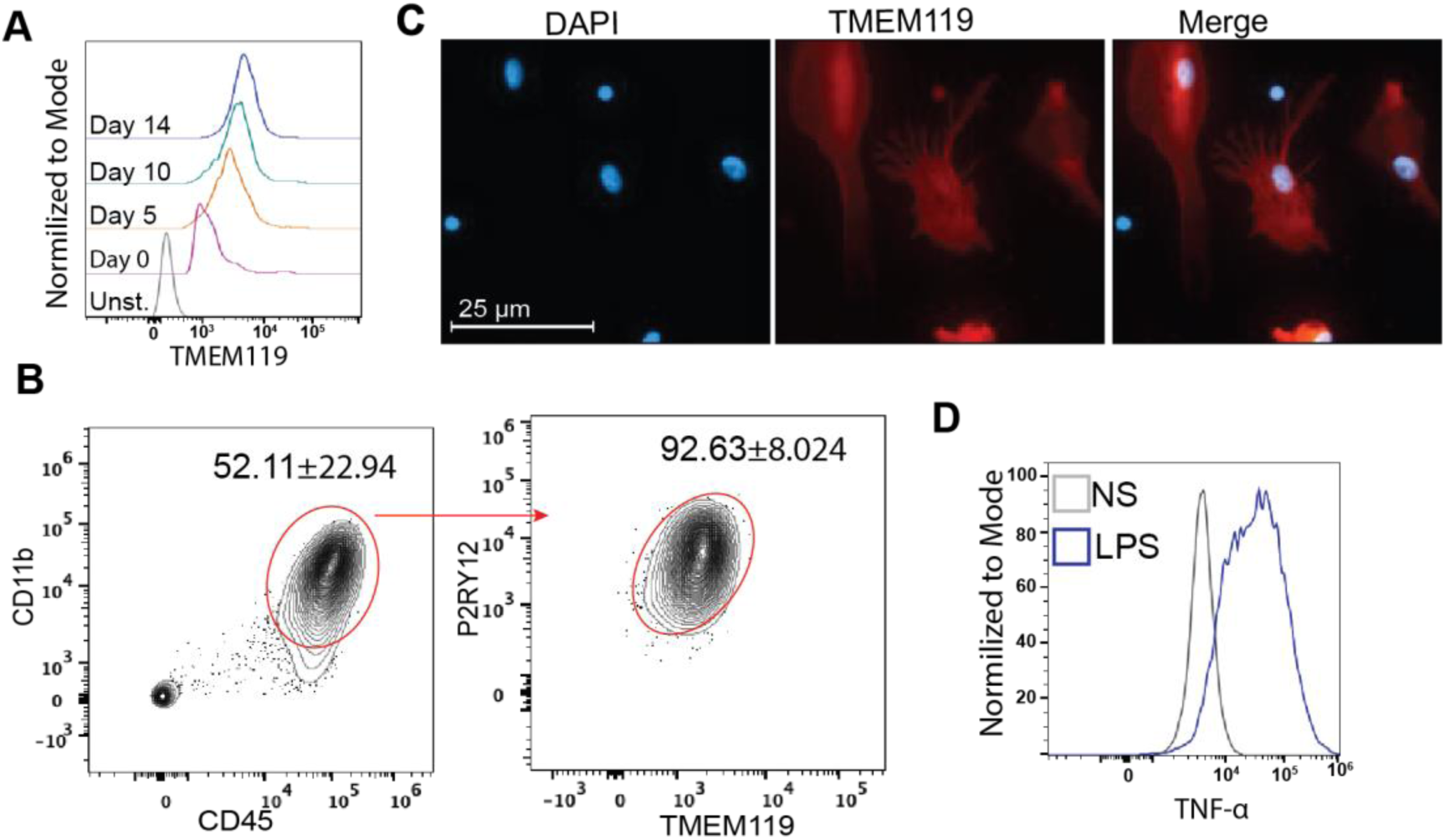
Expression of microglia makers enriched in PBMC-derived microglia. **A)** TMEM119 expression throughout differentiation. **B)** Frequency of CD45^+^CD11b^+^ progenitor cells and P2RY12^+^TMEM119^+^ within this population as measured by flow cytometry. **C)** Confocal imaging of TMEM119 staining of iMGLs. **D)** TNF-⍺ expression by iMGLs before and after LPS was measured by flow cytometry.

**Supp. Fig. 2.**
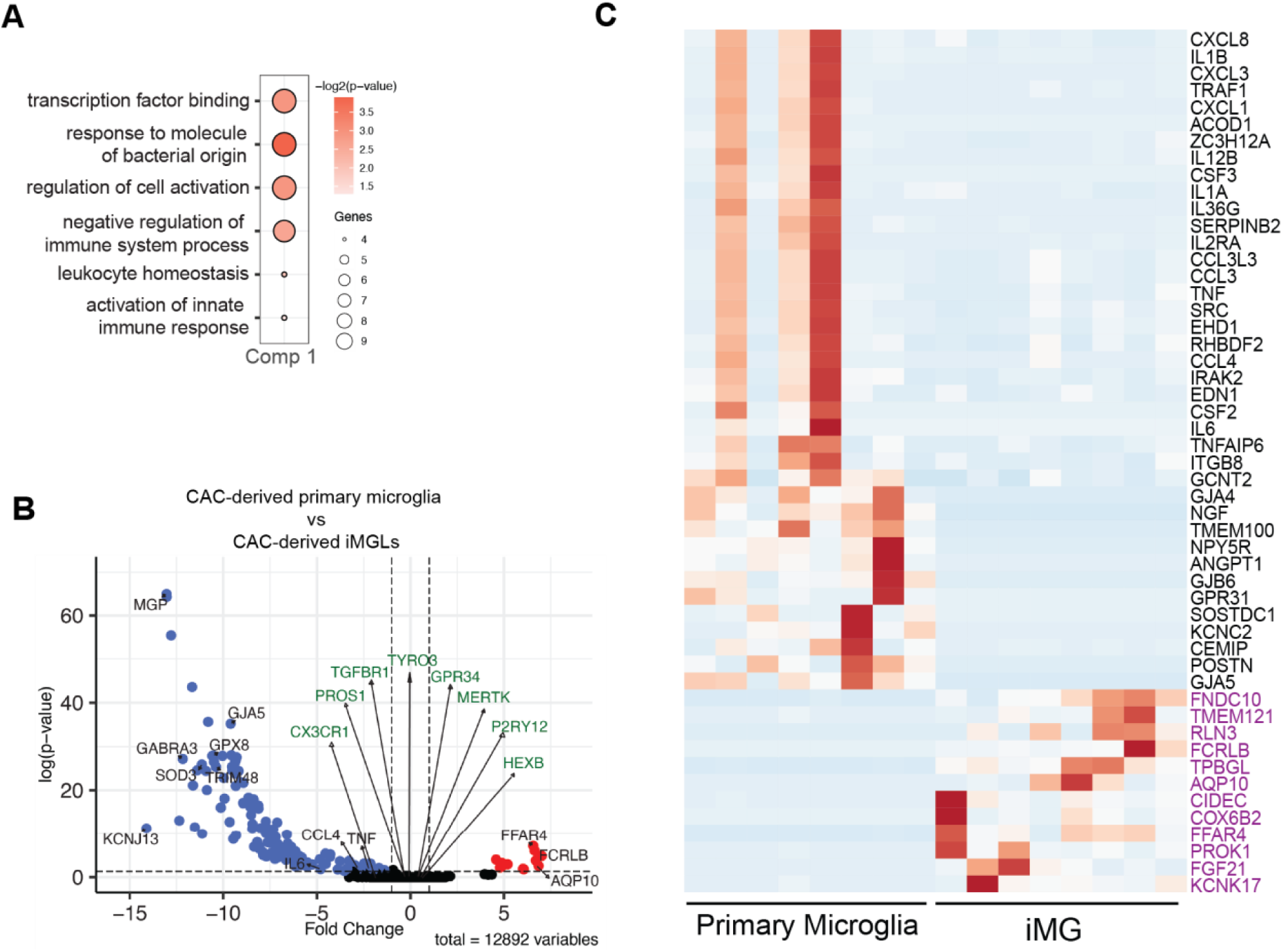
PBMC-derived microglia exhibited transcriptional similarities to primary microglia. **A)** Functional enrichment of genes driving the differences observed in component 1. **B)** Volcano plot depicting DEGs in iMGLs compared to primary microglia. **C)** Heatmap depicts genes mapping to select downregulated GO terms from Fig.1F and all upregulated DEGs (purple).

**Supp. Fig. 3.**
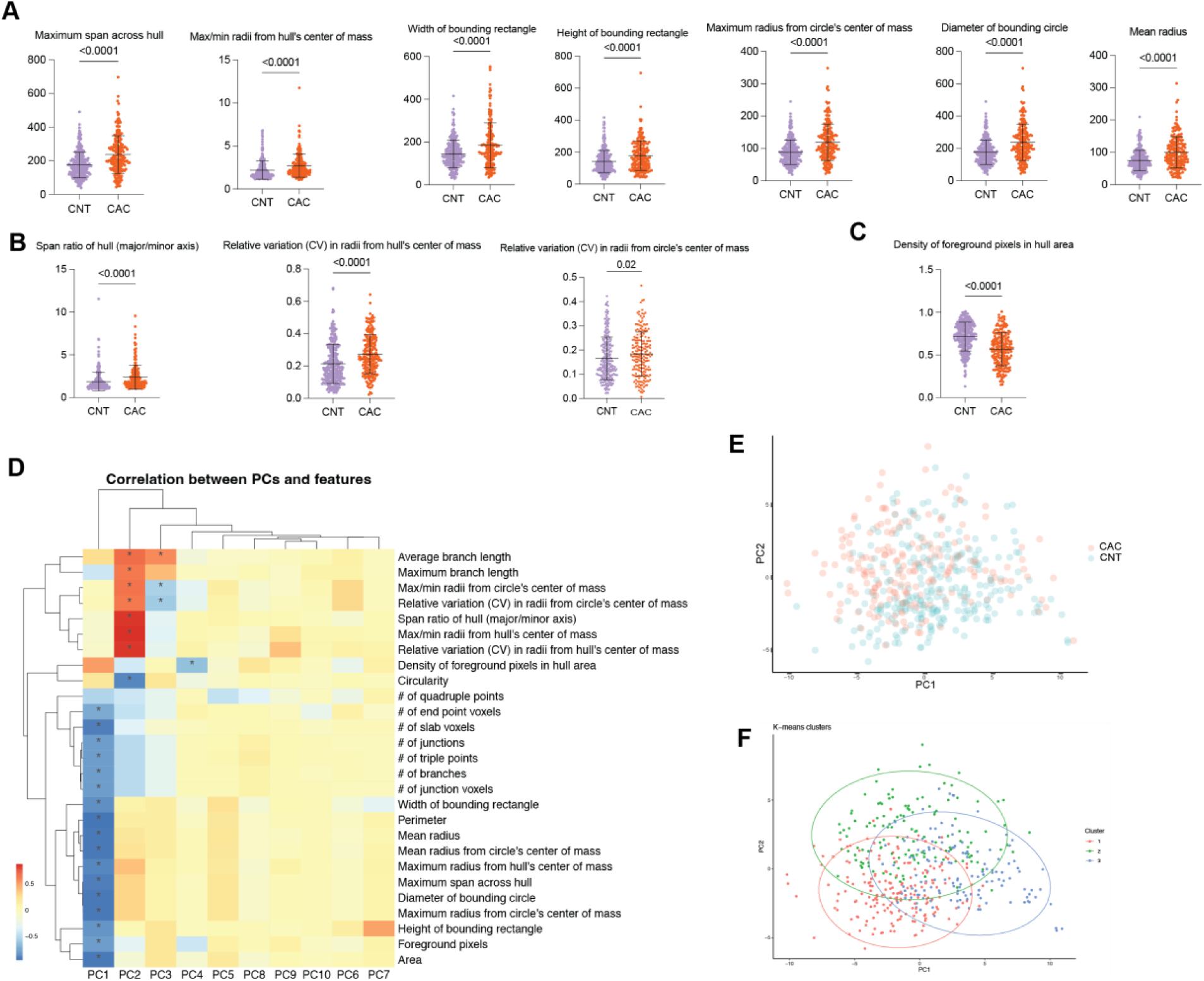
CAC transformed iMGLs morphology. A-C) Morphological measures that were significantly different between control- and CAC-derived iMGLs (p≤0.05). **D)** Spearman’s correlation of 27 morphology features to the first four PCs followed by dimensionality reduction using Principal Components Analysis (PCA) (*abs(R)≥0.55; p≤0.05). **E**) Control and CAC iMGLs were displayed in PCs 1–2 space. **F)** Cells were clustered into three classes using k-means clustering. D-G were generated by MicrogliaMorphologyR package using the default setting unless otherwise specified.

**Supp. Fig. 4.**
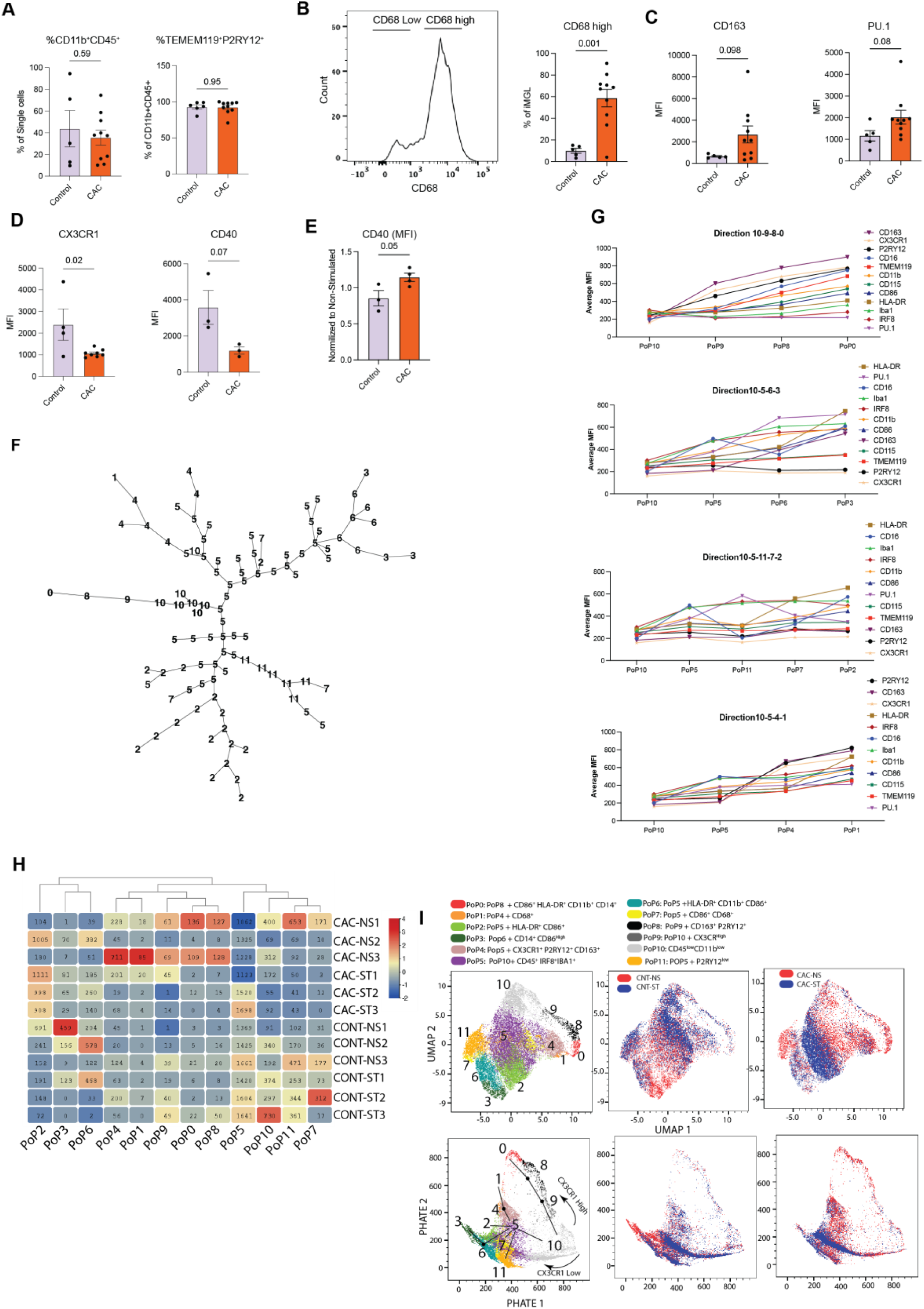
CAC skews phenotype of iMGLs. **A**) The total count of iMGLs and capacity of CD11b^+^CD45^+^ progenitor cells to differentiate to TMEM119^+^P2RY12^+^ cells was determined by flow cytometry. **B**) iMGLs were categorized into CD68^high^ and CD68^low^. The percentage of CD68 high cells increased on CAC-derived iMGLs. **C**) Expression of CD163 and PU.1, **D**) CX3CR1 and CD40 was determined on iMGLs. **E**) Frequency of CD40 and CD68^high^ iMGLs were determined following LPS stimulation. **F**) FlowSOM tree shows the connection of clusters. **G**) Expression of top markers across clusters. **H)** Mean expression of the population in samples. **I**) Both groups and identified clusters were mapped on the UMAP and PHATE plots. Two group comparisons were conducted using either an unpaired t-test with Welch’s correction (for parametric data) or the Mann-Whitney test (for non-parametric data). Error bars in all graphs represent ± standard deviation (SD).

**Supp. Fig. 5.**
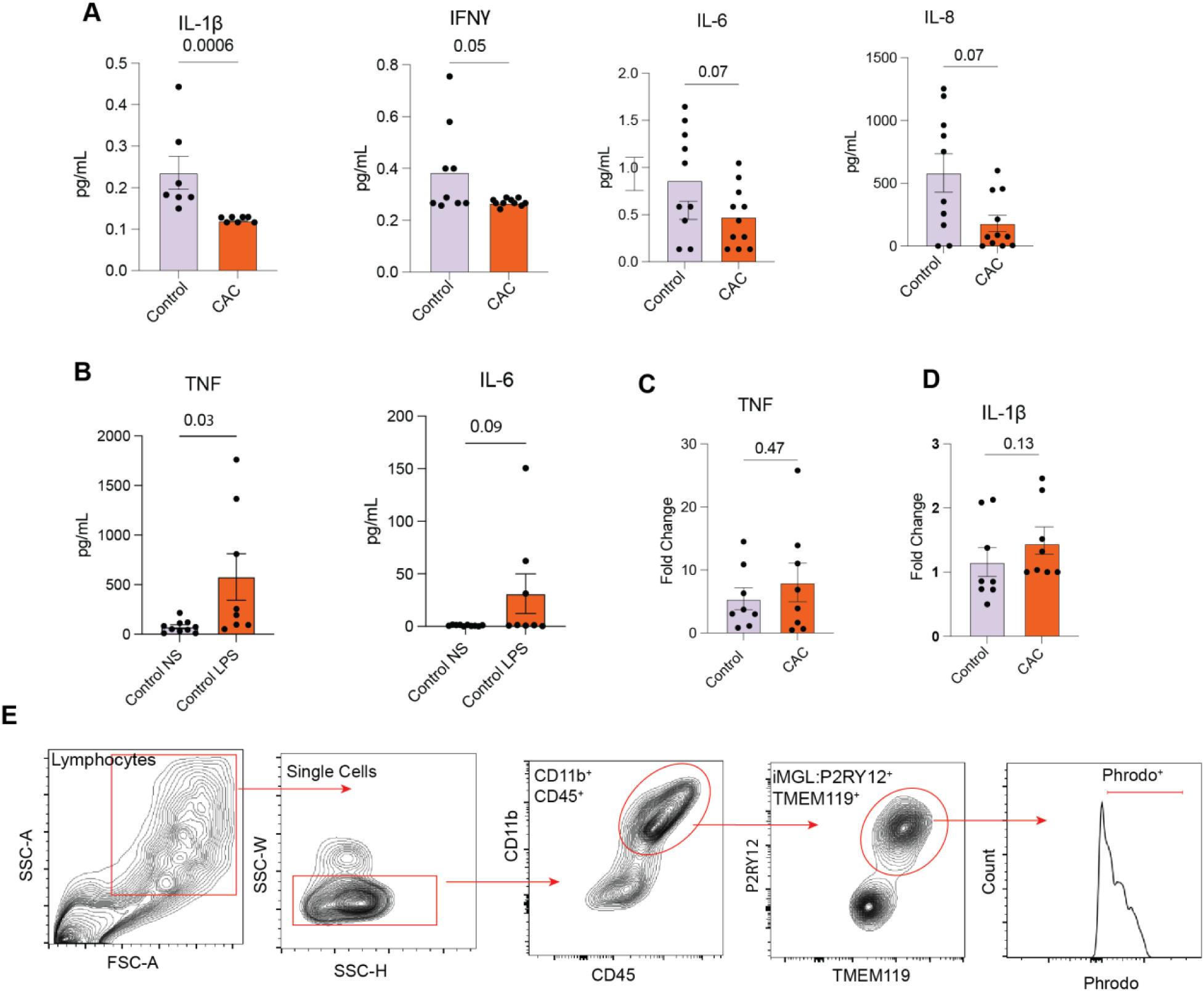
Chronic alcohol consumption alters the function of iMGLs. **A)** Production of IL-1β and IFN-γ, IL-6 and IL-8 at resting, and **B**) TNF and IL-6 in control iMGLs at nonstimulated (NS) and stimulated (LPS) conditions. **C**) Production of TNF and **D)** IL-1β after LPS stimulation. Cytokine expressions were normalized to their unstimulated counterparts. **E)** Gating strategy used to determine phagocytic capacity.

## REFERENCES

1. Delphin-Rittmon ME. The National Survey on Drug Use and Health. 2020. Substance Abuse and Mental Health Services Administration. 2022.

2. SAMHSA CfBHSaQ. National Survey on Drug Use and Health. 2022 [cited 2023 December 18, 2023]. Available from: https://www.samhsa.gov/data/sites/default/files/reports/rpt42728/NSDUHDetailedTabs2022/NSDUHDetailedTabs2022/NSDUHDetTabsSect2pe2022.htm.

3. Smothers BA, Yahr HT. Alcohol use disorder and illicit drug use in admissions to general hospitals in the United States. American Journal on Addictions. 2005;14(3):256–67.

4. Delgado-Rodríguez M, Gómez-Ortega A, Mariscal-Ortiz M, Palma-Pérez S, Sillero-Arenas M. Alcohol drinking as a predictor of intensive care and hospital mortality in general surgery: a prospective study. Addiction. 2003;98(5):611–6.

5. Shield KD, Parry C, Rehm J. Chronic diseases and conditions related to alcohol use. Alcohol research: current reviews. 2014;35(2):155.

6. McIntosh C, Chick J. Alcohol and the nervous system. J Neurol Neurosurg Psychiatry. 2004;75 Suppl 3(Suppl 3):iii16–21. doi: 10.1136/jnnp.2004.045708. PubMed PMID: 15316040; PMCID: PMC1765664.

7. Alekseenko Y. Alcohol-related neurological disorders. In: obert P. Lisak DDT, William M. Carroll and Roongroj Bhidayasiri, editor. International Neurology2016.

8. Hammoud N, Jimenez-Shahed J. Chronic Neurologic Effects of Alcohol. Clin Liver Dis. 2019;23(1):141–55. doi: 10.1016/j.cld.2018.09.010. PubMed PMID: 30454828.

9. Noble JM, Weimer LH. Neurologic complications of alcoholism. Continuum (Minneap Minn). 2014;20(3 Neurology of Systemic Disease):624–41. doi: 10.1212/01.CON.0000450970.99322.84. PubMed PMID: 24893238; PMCID: PMC10563903.

10. Harper C. The Neuropathology of Alcohol-Related Brain Damage. Alcohol and Alcoholism. 2009;44(2):136–40. doi: 10.1093/alcalc/agn102.

11. Sliedrecht W, de Waart R, Witkiewitz K, Roozen HG. Alcohol use disorder relapse factors: A systematic review. Psychiatry Res. 2019;278:97–115. Epub 20190525. doi: 10.1016/j.psychres.2019.05.038. PubMed PMID: 31174033.

12. Bernardin F, Maheut-Bosser A, Paille F. Cognitive impairments in alcohol-dependent subjects. Front Psychiatry. 2014;5:78. Epub 20140716. doi: 10.3389/fpsyt.2014.00078. PubMed PMID: 25076914; PMCID: PMC4099962.

13. Almeida OP, Hankey GJ, Yeap BB, Golledge J, Flicker L. Alcohol consumption and cognitive impairment in older men: a mendelian randomization study. Neurology. 2014;82(12):1038–44.

14. Abrahao KP, Salinas AG, Lovinger DM. Alcohol and the brain: neuronal molecular targets, synapses, and circuits. Neuron. 2017;96(6):1223–38.

15. Egervari G, Siciliano CA, Whiteley EL, Ron D. Alcohol and the brain: from genes to circuits. Trends Neurosci. 2021;44(12):1004–15. Epub 20211023. doi: 10.1016/j.tins.2021.09.006. PubMed PMID: 34702580; PMCID: PMC8616825.

16. Henriques JF, Portugal CC, Canedo T, Relvas JB, Summavielle T, Socodato R. Microglia and alcohol meet at the crossroads: Microglia as critical modulators of alcohol neurotoxicity. Toxicology letters. 2018;283:21–31.

17. Tremblay M-È, Stevens B, Sierra A, Wake H, Bessis A, Nimmerjahn A. The role of microglia in the healthy brain. Journal of Neuroscience. 2011;31(45):16064–9.

18. Tay TL, Savage JC, Hui CW, Bisht K, Tremblay MÈ. Microglia across the lifespan: from origin to function in brain development, plasticity and cognition. The Journal of physiology. 2017;595(6):1929–45.

19. Michell-Robinson MA, Touil H, Healy LM, Owen DR, Durafourt BA, Bar-Or A, Antel JP, Moore CS. Roles of microglia in brain development, tissue maintenance and repair. Brain. 2015;138(5):1138–59.

20. De Sousa RAL, Cassilhas RC. Microglia role as the regulator of cognitive function. Rev Assoc Med Bras (1992). 2023;69(7):e20230412. Epub 20230717. doi: 10.1590/1806-9282.20230412. PubMed PMID: 37466612; PMCID: PMC10352012.

21. Cornell J, Salinas S, Huang HY, Zhou M. Microglia regulation of synaptic plasticity and learning and memory. Neural Regen Res. 2022;17(4):705–16. doi: 10.4103/1673-5374.322423. PubMed PMID: 34472455; PMCID: PMC8530121.

22. Ibáñez C, Acuña T, Quintanilla ME, Pérez-Reytor D, Morales P, Karahanian E. Fenofibrate Decreases Ethanol-Induced Neuroinflammation and Oxidative Stress and Reduces Alcohol Relapse in Rats by a PPAR-α-Dependent Mechanism. Antioxidants. 2023;12(9):1758. doi: 10.3390/antiox12091758.

23. Warden AS, Wolfe SA, Khom S, Varodayan FP, Patel RR, Steinman MQ, Bajo M, Montgomery SE, Vlkolinsky R, Nadav T, Polis I, Roberts AJ, Mayfield RD, Harris RA, Roberto M. Microglia Control Escalation of Drinking in Alcohol-Dependent Mice: Genomic and Synaptic Drivers. Biol Psychiatry. 2020;88(12):910–21. Epub 20200519. doi: 10.1016/j.biopsych.2020.05.011. PubMed PMID: 32680583; PMCID: PMC7674270.

24. Warden AS, Triplett TA, Lyu A, Grantham EK, Azzam MM, DaCosta A, Mason S, Blednov YA, Ehrlich LIR, Mayfield RD, Harris RA. Microglia depletion and alcohol: Transcriptome and behavioral profiles. Addict Biol. 2021;26(2):e12889. Epub 20200316. doi: 10.1111/adb.12889. PubMed PMID: 32176824; PMCID: PMC8510547.

25. Agrawal RG, Hewetson A, George CM, Syapin PJ, Bergeson SE. Minocycline reduces ethanol drinking. Brain Behav Immun. 2011;25 Suppl 1(Suppl 1):S165–9. Epub 20110321. doi: 10.1016/j.bbi.2011.03.002. PubMed PMID: 21397005; PMCID: PMC3098317.

26. Gajbhiye SV, Tripathi RK, Petare A, Potey AV, Shankar A. Minocycline in alcohol withdrawal induced anxiety and alcohol relapse in rats. Current clinical pharmacology. 2018;13(1):65–72.

27. Sabate-Soler S, Bernini M, Schwamborn JC. Immunocompetent brain organoids— microglia enter the stage. Progress in Biomedical Engineering. 2022;4(4). doi: 10.1088/2516-1091/ac8dcf.

28. Speicher AM, Wiendl H, Meuth SG, Pawlowski M. Generating microglia from human pluripotent stem cells: novel in vitro models for the study of neurodegeneration. Molecular Neurodegeneration. 2019;14(1). doi: 10.1186/s13024-019-0347-z.

29. Aktories P, Petry P, Kierdorf K. Microglia in a Dish—Which Techniques Are on the Menu for Functional Studies? Frontiers in Cellular Neuroscience. 2022;16. doi: 10.3389/fncel.2022.908315.

30. Timmerman R, Burm SM, Bajramovic JJ. An Overview of in vitro Methods to Study Microglia. Frontiers in Cellular Neuroscience. 2018;12. doi: 10.3389/fncel.2018.00242.

31. Zhang Y, Cui D. Evolving Models and Tools for Microglial Studies in the Central Nervous System. Neurosci Bull. 2021;37(8):1218–33. Epub 20210609. doi: 10.1007/s12264-021-00706-8. PubMed PMID: 34106404; PMCID: PMC8353053.

32. Warden AS, Han C, Hansen E, Trescott S, Nguyen C, Kim R, Schafer D, Johnson A, Wright M, Ramirez G, Lopez-Sanchez M, Coufal NG. Tools for studying human microglia: In vitro and in vivo strategies. Brain Behav Immun. 2023;107:369–82. Epub 20221103. doi: 10.1016/j.bbi.2022.10.008. PubMed PMID: 36336207; PMCID: PMC9810377.

33. Zhang W, Jiang J, Xu Z, Yan H, Tang B, Liu C, Chen C, Meng Q. Microglia-containing human brain organoids for the study of brain development and pathology. Mol Psychiatry. 2023;28(1):96–107. Epub 20221206. doi: 10.1038/s41380-022-01892-1. PubMed PMID: 36474001; PMCID: PMC9734443.

34. Sheridan SD, Thanos JM, De Guzman RM, McCrea LT, Horng JE, Fu T, Sellgren CM, Perlis RH, Edlow AG. Umbilical cord blood-derived microglia-like cells to model COVID-19 exposure. Transl Psychiatry. 2021;11(1):179. Epub 20210319. doi: 10.1038/s41398-021-01287-w. PubMed PMID: 33741894; PMCID: PMC7976669.

35. Sellgren CM, Gracias J, Watmuff B, Biag JD, Thanos JM, Whittredge PB, Fu T, Worringer K, Brown HE, Wang J, Kaykas A, Karmacharya R, Goold CP, Sheridan SD, Perlis RH. Increased synapse elimination by microglia in schizophrenia patient-derived models of synaptic pruning. Nat Neurosci. 2019;22(3):374–85. Epub 20190204. doi: 10.1038/s41593-018-0334-7. PubMed PMID: 30718903; PMCID: PMC6410571.

36. Sellgren CM, Sheridan SD, Gracias J, Xuan D, Fu T, Perlis RH. Patient-specific models of microglia-mediated engulfment of synapses and neural progenitors. Mol Psychiatry. 2017;22(2):170–7. Epub 20161213. doi: 10.1038/mp.2016.220. PubMed PMID: 27956744; PMCID: PMC5285468.

37. Banerjee A, Lu Y, Do K, Mize T, Wu X, Chen X, Chen J. Validation of Induced Microglia-Like Cells (iMG Cells) for Future Studies of Brain Diseases. Front Cell Neurosci. 2021;15:629279. Epub 20210409. doi: 10.3389/fncel.2021.629279. PubMed PMID: 33897370; PMCID: PMC8063054.

38. Abud EM, Ramirez RN, Martinez ES, Healy LM, Nguyen CHH, Newman SA, Yeromin AV, Scarfone VM, Marsh SE, Fimbres C, Caraway CA, Fote GM, Madany AM, Agrawal A, Kayed R, Gylys KH, Cahalan MD, Cummings BJ, Antel JP, Mortazavi A, Carson MJ, Poon WW, Blurton-Jones M. iPSC-Derived Human Microglia-like Cells to Study Neurological Diseases. Neuron. 2017;94(2):278–93 e9. doi: 10.1016/j.neuron.2017.03.042. PubMed PMID: 28426964; PMCID: PMC5482419.

39. Baker EJ, Farro J, Gonzales S, Helms C, Grant KA. Chronic alcohol self-administration in monkeys shows long-term quantity/frequency categorical stability. Alcohol Clin Exp Res. 2014;38(11):2835–43. doi: 10.1111/acer.12547. PubMed PMID: 25421519; PMCID: PMC4244650.

40. Grant KA, Leng X, Green HL, Szeliga KT, Rogers LS, Gonzales SW. Drinking typography established by scheduled induction predicts chronic heavy drinking in a monkey model of ethanol self-administration. Alcohol Clin Exp Res. 2008;32(10):1824–38. Epub 20080812. doi: 10.1111/j.1530-0277.2008.00765.x. PubMed PMID: 18702645; PMCID: PMC2847427.

41. Jimenez VA, Helms CM, Cornea A, Meshul CK, Grant KA. An ultrastructural analysis of the effects of ethanol self-administration on the hypothalamic paraventricular nucleus in rhesus macaques. Front Cell Neurosci. 2015;9:260. Epub 20150714. doi: 10.3389/fncel.2015.00260. PubMed PMID: 26236193; PMCID: PMC4500925.

42. Grant KA, Leng X, Green HL, Szeliga KT, Rogers LS, Gonzales SW. Drinking typography established by scheduled induction predicts chronic heavy drinking in a monkey model of ethanol self-administration. Alcoholism: Clinical and Experimental Research. 2008;32(10):1824–38.

43. Baker EJ, Farro J, Gonzales S, Helms C, Grant KA. Chronic alcohol self-administration in monkeys shows long-term quantity/frequency categorical stability. Alcoholism: Clinical and Experimental Research. 2014;38(11):2835–43.

44. Bordt EA, Block CL, Petrozziello T, Sadri-Vakili G, Smith CJ, Edlow AG, Bilbo SD. Isolation of Microglia from Mouse or Human Tissue. STAR Protoc. 2020;1(1). Epub 20200603. doi: 10.1016/j.xpro.2020.100035. PubMed PMID: 32783030; PMCID: PMC7416840.

45. Mizee MR, Miedema SS, van der Poel M, Adelia, Schuurman KG, van Strien ME, Melief J, Smolders J, Hendrickx DA, Heutinck KM, Hamann J, Huitinga I. Isolation of primary microglia from the human post-mortem brain: effects of ante- and post-mortem variables. Acta Neuropathol Commun. 2017;5(1):16. Epub 20170217. doi: 10.1186/s40478-017-0418-8. PubMed PMID: 28212663; PMCID: PMC5316206.

46. Lewis SA, Sureshchandra S, Doratt B, Jimenez VA, Stull C, Grant KA, Messaoudi I. Transcriptional, Epigenetic, and Functional Reprogramming of Monocytes From Non-Human Primates Following Chronic Alcohol Drinking. Front Immunol. 2021;12:724015. Epub 20210820. doi: 10.3389/fimmu.2021.724015. PubMed PMID: 34489976; PMCID: PMC8417707.

47. Kim D, Langmead B, Salzberg SL. HISAT: a fast spliced aligner with low memory requirements. Nat Methods. 2015;12(4):357–60. Epub 20150309. doi: 10.1038/nmeth.3317. PubMed PMID: 25751142; PMCID: PMC4655817.

48. Kinsella RJ, Kahari A, Haider S, Zamora J, Proctor G, Spudich G, Almeida-King J, Staines D, Derwent P, Kerhornou A, Kersey P, Flicek P. Ensembl BioMarts: a hub for data retrieval across taxonomic space. Database (Oxford). 2011;2011:bar030. Epub 20110723. doi: 10.1093/database/bar030. PubMed PMID: 21785142; PMCID: PMC3170168.

49. Rhoades NS, Hendrickson SM, Prongay K, Haertel A, Gill L, Edwards RA, Garzel L, Slifka MK, Messaoudi I. Growth faltering regardless of chronic diarrhea is associated with mucosal immune dysfunction and microbial dysbiosis in the gut lumen. Mucosal Immunol. 2021;14(5):1113–26. Epub 20210622. doi: 10.1038/s41385-021-00418-2. PubMed PMID: 34158595; PMCID: PMC8379072.

50. Rhoades N, Barr T, Hendrickson S, Prongay K, Haertel A, Gill L, Garzel L, Whiteson K, Slifka M, Messaoudi I. Maturation of the infant rhesus macaque gut microbiome and its role in the development of diarrheal disease. Genome Biology. 2019;20(1). doi: 10.1186/s13059-019-1789-x.

51. Zhou Y, Zhou B, Pache L, Chang M, Khodabakhshi AH, Tanaseichuk O, Benner C, Chanda SK. Metascape provides a biologist-oriented resource for the analysis of systems-level datasets. Nat Commun. 2019;10(1):1523. Epub 20190403. doi: 10.1038/s41467-019-09234-6. PubMed PMID: 30944313; PMCID: PMC6447622.

52. Subramanian A, Tamayo P, Mootha VK, Mukherjee S, Ebert BL, Gillette MA, Paulovich A, Pomeroy SL, Golub TR, Lander ES, Mesirov JP. Gene set enrichment analysis: a knowledge-based approach for interpreting genome-wide expression profiles. Proc Natl Acad Sci U S A. 2005;102(43):15545–50. Epub 20050930. doi: 10.1073/pnas.0506580102. PubMed PMID: 16199517; PMCID: PMC1239896.

53. Kim J, Pavlidis P, Ciernia AV. Development of a High-Throughput Pipeline to Characterize Microglia Morphological States at a Single-Cell Resolution. eNeuro. 2024;11(7). Epub 20240730. doi: 10.1523/ENEURO.0014-24.2024. PubMed PMID: 39029952; PMCID: PMC11289588.

54. Korzhevskii DE, Kirik OV. Brain Microglia and Microglial Markers. Neuroscience and Behavioral Physiology. 2016;46(3):284-90. doi: 10.1007/s11055-016-0231-z.

55. Böttcher C, Schlickeiser S, Sneeboer MAM, Kunkel D, Knop A, Paza E, Fidzinski P, Kraus L, Snijders GJL, Kahn RS, Schulz AR, Mei HE, Hol EM, Siegmund B, Glauben R, Spruth EJ, De Witte LD, Priller J. Human microglia regional heterogeneity and phenotypes determined by multiplexed single-cell mass cytometry. Nature Neuroscience. 2019;22(1):78–90. doi: 10.1038/s41593-018-0290-2.

56. Monaco G, Chen H, Poidinger M, Chen J, de Magalhaes JP, Larbi A. flowAI: automatic and interactive anomaly discerning tools for flow cytometry data. Bioinformatics. 2016;32(16):2473–80. Epub 20160410. doi: 10.1093/bioinformatics/btw191. PubMed PMID: 27153628.

57. Pedersen CB, Dam SH, Barnkob MB, Leipold MD, Purroy N, Rassenti LZ, Kipps TJ, Nguyen J, Lederer JA, Gohil SH, Wu CJ, Olsen LR. cyCombine allows for robust integration of single-cell cytometry datasets within and across technologies. Nat Commun. 2022;13(1):1698. Epub 20220331. doi: 10.1038/s41467-022-29383-5. PubMed PMID: 35361793; PMCID: PMC8971492.

58. McInnes L, Healy J, Saul N, Großberger L. UMAP: Uniform Manifold Approximation and Projection. Journal of Open Source Software. 2018;3(29). doi: 10.21105/joss.00861.

59. Kratochvil M, Koladiya A, Vondrasek J. Generalized EmbedSOM on quadtree-structured self-organizing maps. F1000Res. 2019;8:2120. Epub 20191218. doi: 10.12688/f1000research.21642.2. PubMed PMID: 32518625; PMCID: PMC7255855.

60. Moon KR, van Dijk D, Wang Z, Gigante S, Burkhardt DB, Chen WS, Yim K, Elzen AVD, Hirn MJ, Coifman RR, Ivanova NB, Wolf G, Krishnaswamy S. Visualizing structure and transitions in high-dimensional biological data. Nat Biotechnol. 2019;37(12):1482–92. Epub 20191203. doi: 10.1038/s41587-019-0336-3. PubMed PMID: 31796933; PMCID: PMC7073148.

61. Kierdorf K, Erny D, Goldmann T, Sander V, Schulz C, Perdiguero EG, Wieghofer P, Heinrich A, Riemke P, Holscher C, Muller DN, Luckow B, Brocker T, Debowski K, Fritz G, Opdenakker G, Diefenbach A, Biber K, Heikenwalder M, Geissmann F, Rosenbauer F, Prinz M. Microglia emerge from erythromyeloid precursors via Pu.1- and Irf8-dependent pathways. Nat Neurosci. 2013;16(3):273–80. Epub 20130120. doi: 10.1038/nn.3318. PubMed PMID: 23334579.

62. Winter A, McMurray KMJ, Ahlbrand R, Allgire E, Shukla S, Jones J, Sah R. The subfornical organ regulates acidosis-evoked fear by engaging microglial acid-sensor TDAG8 and forebrain neurocircuits in male mice. J Neurosci Res. 2022;100(9):1732–46. Epub 20220512. doi: 10.1002/jnr.25059. PubMed PMID: 35553084; PMCID: PMC9812228.

63. Campbell EJ, Lawrence AJ. It’s more than just interoception: The insular cortex involvement in alcohol use disorder. J Neurochem. 2021;157(5):1644–51. Epub 20210210. doi: 10.1111/jnc.15310. PubMed PMID: 33486788.

64. Hitzemann R, Gao L, Fei SS, Ray K, Vigh-Conrad KA, Phillips TJ, Searles R, Cervera-Juanes RP, Khadka R, Carlson TL, Gonzales SW, Newman N, Grant KA. Effects of repeated alcohol abstinence on within-subject prefrontal cortical gene expression in rhesus macaques. Adv Drug Alcohol Res. 2024;4:12528. Epub 20240426. doi: 10.3389/adar.2024.12528. PubMed PMID: 38737578; PMCID: PMC11082748.

65. Walter TJ, Vetreno RP, Crews FT. Alcohol and Stress Activation of Microglia and Neurons: Brain Regional Effects. Alcohol Clin Exp Res. 2017;41(12):2066–81. Epub 20171108. doi: 10.1111/acer.13511. PubMed PMID: 28941277; PMCID: PMC5725687.

66. Lier J, Streit WJ, Bechmann I. Beyond Activation: Characterizing Microglial Functional Phenotypes. Cells. 2021;10(9). Epub 20210828. doi: 10.3390/cells10092236. PubMed PMID: 34571885; PMCID: PMC8464670.

67. Bohlen CJ, Bennett FC, Tucker AF, Collins HY, Mulinyawe SB, Barres BA. Diverse Requirements for Microglial Survival, Specification, and Function Revealed by Defined-Medium Cultures. Neuron. 2017;94(4):759–73 e8. doi: 10.1016/j.neuron.2017.04.043. PubMed PMID: 28521131; PMCID: PMC5523817.

68. Tewari M, Khan M, Verma M, Coppens J, Kemp JM, Bucholz R, Mercier P, Egan TM. Physiology of Cultured Human Microglia Maintained in a Defined Culture Medium. Immunohorizons. 2021;5(4):257–72. Epub 20210430. doi: 10.4049/immunohorizons.2000101. PubMed PMID: 33931497; PMCID: PMC9190148.

69. Stratoulias V, Venero JL, Tremblay ME, Joseph B. Microglial subtypes: diversity within the microglial community. EMBO J. 2019;38(17):e101997. Epub 20190802. doi: 10.15252/embj.2019101997. PubMed PMID: 31373067; PMCID: PMC6717890.

70. Lauro C, Limatola C. Metabolic Reprograming of Microglia in the Regulation of the Innate Inflammatory Response. Front Immunol. 2020;11:493. Epub 20200320. doi: 10.3389/fimmu.2020.00493. PubMed PMID: 32265936; PMCID: PMC7099404.

71. Yu T, Zhang X, Shi H, Tian J, Sun L, Hu X, Cui W, Du D. P2Y12 regulates microglia activation and excitatory synaptic transmission in spinal lamina II neurons during neuropathic pain in rodents. Cell Death Dis. 2019;10(3):165. Epub 20190218. doi: 10.1038/s41419-019-1425-4. PubMed PMID: 30778044; PMCID: PMC6379416.

72. Ruan C, Elyaman W. A New Understanding of TMEM119 as a Marker of Microglia. Front Cell Neurosci. 2022;16:902372. Epub 20220613. doi: 10.3389/fncel.2022.902372. PubMed PMID: 35769325; PMCID: PMC9234454.

73. Janda E, Boi L, Carta AR. Microglial Phagocytosis and Its Regulation: A Therapeutic Target in Parkinson’s Disease? Front Mol Neurosci. 2018;11:144. Epub 20180427. doi: 10.3389/fnmol.2018.00144. PubMed PMID: 29755317; PMCID: PMC5934476.

74. Smith AM, Gibbons HM, Oldfield RL, Bergin PM, Mee EW, Faull RL, Dragunow M. The transcription factor PU.1 is critical for viability and function of human brain microglia. Glia. 2013;61(6):929–42. Epub 20130309. doi: 10.1002/glia.22486. PubMed PMID: 23483680.

75. Lee W, Kim HS, Baek SY, Lee GR. Transcription factor IRF8 controls Th1-like regulatory T-cell function. Cell Mol Immunol. 2016;13(6):785–94. Epub 20150713. doi: 10.1038/cmi.2015.72. PubMed PMID: 26166768; PMCID: PMC5101446.

76. Masuda T, Tsuda M, Yoshinaga R, Tozaki-Saitoh H, Ozato K, Tamura T, Inoue K. IRF8 is a critical transcription factor for transforming microglia into a reactive phenotype. Cell Rep. 2012;1(4):334–40. Epub 20120405. doi: 10.1016/j.celrep.2012.02.014. PubMed PMID: 22832225; PMCID: PMC4158926.

77. Nixon SJ, Prather R, Lewis B. Sex differences in alcohol-related neurobehavioral consequences. Handb Clin Neurol. 2014;125:253–72. doi: 10.1016/B978-0-444-62619-6.00016-1. PubMed PMID: 25307580.

78. Petry NM. Delay discounting of money and alcohol in actively using alcoholics, currently abstinent alcoholics, and controls. Psychopharmacology (Berl). 2001;154(3):243–50. doi: 10.1007/s002130000638. PubMed PMID: 11351931.

79. Melbourne JK, Thompson KR, Peng H, Nixon K. Its complicated: The relationship between alcohol and microglia in the search for novel pharmacotherapeutic targets for alcohol use disorders. Prog Mol Biol Transl Sci. 2019;167:179–221. Epub 20190729. doi: 10.1016/bs.pmbts.2019.06.011. PubMed PMID: 31601404.

80. Henriques JF, Portugal CC, Canedo T, Relvas JB, Summavielle T, Socodato R. Microglia and alcohol meet at the crossroads: Microglia as critical modulators of alcohol neurotoxicity. Toxicol Lett. 2018;283:21–31. Epub 20171110. doi: 10.1016/j.toxlet.2017.11.002. PubMed PMID: 29129797.

81. Leng F, Edison P. Neuroinflammation and microglial activation in Alzheimer disease: where do we go from here? Nat Rev Neurol. 2021;17(3):157–72. Epub 20201214. doi: 10.1038/s41582-020-00435-y. PubMed PMID: 33318676.

82. Lan L, Wang H, Zhang X, Shen Q, Li X, He L, Rong X, Peng J, Mo J, Peng Y. Chronic exposure of alcohol triggers microglia-mediated synaptic elimination inducing cognitive impairment. Exp Neurol. 2022;353:114061. Epub 20220401. doi: 10.1016/j.expneurol.2022.114061. PubMed PMID: 35367455.

83. Sureshchandra S, Stull C, Ligh BJK, Nguyen SB, Grant KA, Messaoudi I. Chronic heavy drinking drives distinct transcriptional and epigenetic changes in splenic macrophages. EBioMedicine. 2019;43:594–606. Epub 20190418. doi: 10.1016/j.ebiom.2019.04.027. PubMed PMID: 31005514; PMCID: PMC6557917.

84. Lewis SA, Doratt BM, Sureshchandra S, Jankeel A, Newman N, Shen W, Grant KA, Messaoudi I. Ethanol Consumption Induces Nonspecific Inflammation and Functional Defects in Alveolar Macrophages. Am J Respir Cell Mol Biol. 2022;67(1):112–24. doi: 10.1165/rcmb.2021-0346OC. PubMed PMID: 35380939; PMCID: PMC9273227.

85. Wu M-Y, Liu L, Wang E-J, Xiao H-T, Cai C-Z, Wang J, Su H, Wang Y, Tan J, Zhang Z, Wang J, Yao M, Ouyang D-F, Yue Z, Li M, Chen Y, Bian Z-X, Lu J-H. PI3KC3 complex subunit NRBF2 is required for apoptotic cell clearance to restrict intestinal inflammation. Autophagy. 2021;17(5):1096–111. doi: 10.1080/15548627.2020.1741332.

86. Li Y, Li J, Yu Q, Ji L, Peng B. METTL14 regulates microglia/macrophage polarization and NLRP3 inflammasome activation after ischemic stroke by the KAT3B-STING axis. Neurobiol Dis. 2023;185:106253. Epub 20230802. doi: 10.1016/j.nbd.2023.106253. PubMed PMID: 37541353.

87. Schmidt C, Schneble N, Muller JP, Bauer R, Perino A, Marone R, Rybalkin SD, Wymann MP, Hirsch E, Wetzker R. Phosphoinositide 3-kinase gamma mediates microglial phagocytosis via lipid kinase-independent control of cAMP. Neuroscience. 2013;233:44–53. Epub 20121229. doi: 10.1016/j.neuroscience.2012.12.036. PubMed PMID: 23276671.

88. Niedzielska M, Bodendorfer B, Munch S, Eichner A, Derigs M, da Costa O, Schweizer A, Neff F, Nitschke L, Sparwasser T, Keyse SM, Lang R. Gene trap mice reveal an essential function of dual specificity phosphatase Dusp16/MKP-7 in perinatal survival and regulation of Toll-like receptor (TLR)-induced cytokine production. J Biol Chem. 2014;289(4):2112–26. Epub 20131205. doi: 10.1074/jbc.M113.535245. PubMed PMID: 24311790; PMCID: PMC3900958.

89. Chen Y, Wang J, Wang X, Li X, Song J, Fang J, Liu X, Liu T, Wang D, Li Q, Wen S, Ma D, Xia J, Luo L, Zheng SG, Cui J, Zeng G, Chen L, Cheng B, Wang Z. Pik3ip1 Is a Negative Immune Regulator that Inhibits Antitumor T-Cell Immunity. Clin Cancer Res. 2019;25(20):6180–94. Epub 20190726. doi: 10.1158/1078-0432.CCR-18-4134. PubMed PMID: 31350312.

90. Monteleone G, Pallone F, MacDonald TT. Smad7 in TGF-beta-mediated negative regulation of gut inflammation. Trends Immunol. 2004;25(10):513–7. doi: 10.1016/j.it.2004.07.008. PubMed PMID: 15364052.

91. Arafeh R, Di Pizio A, Elkahloun AG, Dym O, Niv MY, Samuels Y. RASA2 and NF1; two-negative regulators of Ras with complementary functions in melanoma. Oncogene. 2019;38(13):2432–4. Epub 20181126. doi: 10.1038/s41388-018-0578-4. PubMed PMID: 30478445.

92. Munoz EM. Microglia in Circumventricular Organs: The Pineal Gland Example. ASN Neuro. 2022;14:17590914221135697. doi: 10.1177/17590914221135697. PubMed PMID: 36317305; PMCID: PMC9629557.

93. Popiolek-Barczyk K, Ciechanowska A, Ciapala K, Pawlik K, Oggioni M, Mercurio D, De Simoni MG, Mika J. The CCL2/CCL7/CCL12/CCR2 pathway is substantially and persistently upregulated in mice after traumatic brain injury, and CCL2 modulates the complement system in microglia. Mol Cell Probes. 2020;54:101671. Epub 20201104. doi: 10.1016/j.mcp.2020.101671. PubMed PMID: 33160071.

94. Morrison H, Young K, Qureshi M, Rowe RK, Lifshitz J. Quantitative microglia analyses reveal diverse morphologic responses in the rat cortex after diffuse brain injury. Sci Rep. 2017;7(1):13211. Epub 20171016. doi: 10.1038/s41598-017-13581-z. PubMed PMID: 29038483; PMCID: PMC5643511.

95. Akhter R, Shao Y, Formica S, Khrestian M, Bekris LM. TREM2 alters the phagocytic, apoptotic and inflammatory response to Abeta(42) in HMC3 cells. Mol Immunol. 2021;131:171–9. Epub 20210115. doi: 10.1016/j.molimm.2020.12.035. PubMed PMID: 33461764; PMCID: PMC8147571.

96. Thuong NT, Tram TT, Dinh TD, Thai PV, Heemskerk D, Bang ND, Chau TT, Russell DG, Thwaites GE, Hawn TR, Caws M, Dunstan SJ. MARCO variants are associated with phagocytosis, pulmonary tuberculosis susceptibility and Beijing lineage. Genes Immun. 2016;17(7):419–25. Epub 20161117. doi: 10.1038/gene.2016.43. PubMed PMID: 27853145; PMCID: PMC5133378.

